# H3K27M diffuse midline glioma is homologous recombination defective and sensitized to radiotherapy and NK cell-mediated antitumor immunity by PARP inhibition

**DOI:** 10.1101/2024.08.26.609803

**Authors:** Yupei Guo, Zian Li, Leslie A. Parsels, Zhuwen Wang, Joshua D. Parsels, Anushka Dalvi, Stephanie The, Nan Hu, Victoria M. Valvo, Robert Doherty, Erik Peterson, Xinjun Wang, Sujatha Venkataraman, Sameer Agnihotri, Sriram Venneti, Daniel R. Wahl, Michael D. Green, Theodore S. Lawrence, Carl Koschmann, Meredith A. Morgan, Qiang Zhang

## Abstract

**Background:** Radiotherapy (RT) is the primary treatment for diffuse midline glioma (DMG), a lethal pediatric malignancy defined by histone H3 lysine 27-to-methionine (H3K27M) mutation. Based on the loss of H3K27 trimethylation producing broad epigenomic alterations, we hypothesized that H3K27M causes a functional double-strand break (DSB) repair defect that could be leveraged therapeutically with PARP inhibitor and RT for selective radiosensitization and antitumor immune responses.

**Methods:** H3K27M isogenic DMG cells and orthotopic brainstem DMG tumors in immune deficient and syngeneic, immune competent mice were used to evaluate the efficacy and mechanisms of PARP1/2 inhibition by olaparib or PARP1 inhibition by AZD9574 with concurrent RT.

**Results:** H3K27M mutation caused an HRR defect characterized by impaired RT-induced K63-linked polyubiquitination of histone H1 and inhibition of HRR protein recruitment. H3K27M DMG cells were selectively radiosensitized by olaparib in comparison to isogenic controls, and this effect translated to efficacy in H3K27M orthotopic brainstem tumors. Olaparib and RT induced an innate immune response and induction of NK cell (NKG2D) activating ligands leading to increased NK cell-mediated lysis of DMG tumor cells. In immunocompetent syngeneic orthotopic DMG tumors, either olaparib or AZD9574 in combination with RT enhanced intratumoral NK cell infiltration and activity in association with NK cell-mediated therapeutic responses and favorable activity of AZD9574.

**Conclusions:** The HRR deficiency in H3K27M DMG can be therapeutically leveraged with PARP inhibitors to radiosensitize and induce an NK cell-mediated antitumor immune response selectively in H3K27M DMG, supporting the clinical investigation of best-in-class PARP inhibitors with RT in DMG patients.

**Key points:**

- H3K27M DMG are HRR defective and selectively radiosensitized by PARP inhibitor.
- PARP inhibitor with RT enhances NKG2D ligand expression and NK cell-mediated lysis.
- NK cells are required for the therapeutic efficacy of PARP inhibitor and RT.

**Importance of the Study:** Radiotherapy is the cornerstone of H3K27M-mutant diffuse midline glioma treatment, but almost all patients succumb to tumor recurrence with poor overall survival, underscoring the need for RT-based precision combination therapy. Here, we reveal HRR deficiency as an H3K27M-mediated vulnerability and identify a novel mechanism linking impaired RT-induced histone H1 polyubiquitination and the subsequent RNF168/BRCA1/RAD51 recruitment in H3K27M DMG. This model is supported by selective radiosensitization of H3K27M DMG by PARP inhibitor. Notably, the combination treatment results in NKG2D ligand expression that confers susceptibility to NK cell killing in H3K27M DMG. We also show that the novel brain penetrant, PARP1-selective inhibitor AZD9574 compares favorably to olaparib when combined with RT, prolonging survival in a syngeneic orthotopic model of H3K27M DMG. This study highlights the ability of PARP1 inhibition to radiosensitize and induce an NK cell-mediated antitumor immunity in H3K27M DMG and supports future clinical investigation.

## Introduction

DMG H3K27-altered is a universally fatal, high grade pediatric brain tumor defined by histone H3 lysine 27-to-methionine (H3K27M) mutation [1, 2]. Given the critical functions of the brainstem and the microscopic extension of tumor into surrounding normal tissue, surgery is not a treatment option and chemotherapeutic drugs have limited efficacy [3, 4]. Fractionated radiotherapy (RT) is the only standard treatment that improves symptoms and delays tumor progression, however, DMGs inevitably recur within the high dose radiation field [5–7]. Characterization of both the H3K27M-mediated cell vulnerabilities and their biological interaction with RT in DMG will inform more effective RT combination treatment options.

H3K27M in DMG is a gain-of-function mutation that inhibits polycomb repressive complex 2 (PRC2) methyltransferase activity, thus leading to drastic epigenetic alteration in both a cis– and trans-manner with global loss of the repressive tri-methylation (H3K27me3) mark and concomitant gain of activating acetylation (H3K27ac) of H3K27 [8–11]. Radiation further alters the epigenome through histone posttranslational modifications, such as ubiquitination and methylation, that are required for the recruitment of DNA repair proteins [12, 13]. Siddaway et al. reported that H3K27M alters DNA repair protein binding, but the unique molecular interaction between RT– and H3K27M-induced epigenetic alterations remains elusive [14]. Evidence supports an interaction between RT and H3K27M in that pharmacological inhibition of the H3 demethylase, JMJD3/UTX, or bromodomain protein 4 (BRD4) sensitized to RT by restoring H3K27me3 and preventing the binding of BRD4 to H3K27ac, respectively [15–17]. Together with increased H3K27ac, H3K27M mutation also causes enhanced acetylation of lysine 16 on histone H4 (H4K16ac) and lysine 9 and 64 on histone H3 (H3K64ac, H3K9ac) [18, 19]. Collectively, these alterations suggest a more hyperacetylated and ‘‘open’’ chromatin state that may disrupt the binding affinity of linker histone H1 to chromatin [20–22]. It is currently unclear how H3K27M affects RT-induced DNA repair through histone H1 in DMG.

In addition to inducing lethal DNA damage in tumor cells, RT also promotes an antitumor immune response [23–26]. Radiotherapy-induced DNA damage increases tumor antigenicity, mitigates immunosuppression, and generates both innate and adaptive antitumor immune responses. Further, we and others have shown that inhibitors of the DNA damage response (DDR) enhance immunogenic DNA/RNA following RT and antitumor immunity via type I interferon (T1IFN) [27–30]. In addition, the combination of DDR inhibition with RT further primes and activates antitumor immune cells such as cytotoxic T and natural killer (NK) cells in the tumor microenvironment (TME) [31–33]. The immune-naïve tumor microenvironment and limited lymphocytes in DMG present challenges in developing effective immunotherapies [34–36]. Despite its immunologically inert nature, DMG is not completely exempt from intratumoral lymphocytes as in vitro and in vivo tumor cell killing assays demonstrated that NK cells elicit more robust cytotoxicity toward DMG tumor cells relative to CD8^+^ T cells [34, 37]. Meanwhile, intratumoral NK cells, but not CD8^+^ T cells, positively correlate with overall survival in H3K27M DMG patients [37] supporting investigation of therapeutic strategies to enhance the NK cell antitumor immune response in H3K27M DMG.

NK cells have direct cytotoxicity against tumor cells through production of granzymes, perforin, and cytokines [38]. Both activating and inhibitory receptors expressed on the surface of NK cells contribute to the execution of NK cell functions [38, 39]. The NK cell surface activating receptors, especially NKG2D, recognize ligands expressed on the surface of cancer cells that when engaged by respective ligands (MICA, MICB, and ULBP1-6 in human; Rae, H60, and Mult1 in mouse) induce degranulation, cytokine production, and NK cell lysis activity [40]. DNA damage induced by RT or genotoxic agents is a potent stimulator of NKG2D ligand expression [41, 42]. Thus, further potentiation of RT-mediated NKG2D ligand presentation may shift the balance of DMG tumor cells toward activation of an NK cell-mediated antitumor immune response.

In this study, we initially found that H3K27M mutation caused a deficiency in homologous recombination repair (HRR) that was associated with impairment of the interaction between linker histone H1 and hyperacetylated core histones and consequently inhibition of RT-induced H1 Lys63 (K63)-linked polyubiquitination and subsequent recruitment of HRR proteins (RNF168, BRCA1, RAD51). These finding led us to hypothesize that the HRR deficiency in H3K27M DMG could be therapeutically leveraged by PARP inhibition for both radiosensitization and enhanced RT-induced immunogenicity. To test this hypothesis, we utilized the FDA approved PARP1/2 inhibitor olaparib as well as the novel blood-brain barrier penetrant PARP1 selective inhibitor AZD9574 [43] in H3K27 isogenic DMG cell lines and immune competent, syngeneic, orthotopic mouse models of H3K27M mutant DMG. Taken together, this work highlights the ability of PARP inhibition including a best-in-class PARP1 selective inhibitor to enhance both radiation sensitivity as well as radiation-induced NK cell-mediated antitumor immune responses in H3K27M mutant DMG.

## Materials and Methods

### Cell culture

Patient-derived H3K27M isogenic (H3K27M and H3K27M-KO) cell line pairs (BT245 and DIPG XIII) were provided by Dr. Nada Jabado (McGill University) [44]. Patient-derived H3K27M-isogenic (H3K27M and H3K27M-knockdown (KD)) cell line pairs (DIPG007) were from Dr. Sriram Venneti at University of Michigan. All human DIPG cell lines were cultured in the tumor stem media (TSM) basic medium supplied with B27 minus vitamin A (Thermo Fisher, #12587010), human recombinant EGF (PeproTech, #AF-100-15), human recombinant bFGF (#100-18B), human recombinant PDGF-AA (#100-13A), human recombinant PDGF-BB (#100-14B), and Heparin (Stem Cell Technologies, #07980). H3K27MPP murine DMG cells were provided by Dr. Sameer Agnihotri (University of Pittsburg) and grown in neurobasal media with complete NeuroCult Proliferation kit (Stem Cell Technologies, #05702), supplemented with EGF, bFGF and heparin [45]. Murine PPK/PPW cells were established from a tumor derived by in utero electroporation of plasmids expressing dominant negative TP53, mutant PDGFRA (D842V), and mutant H3K27M or H3WT, respectively [46]. U2OS (DR-GFP), 293T and Hela cells were grown in DMEM with 10% FBS, 1% penicillin-streptomycin.

### In vivo orthotopic DMG model and treatment

The animal studies were approved by University of Michigan Institutional Animal Care & Use Committee. An implantable guide-screw system was used to generate orthotopic brainstem DMG models as previously described [47]. PARP inhibitors olaparib and AZD9574 were provided by AstraZeneca. Olaparib was dissolved in DMSO and then diluted in 10% 2-hydroxypropyl-β-cyclodextrin (Cayman Chemicals), and AZD9574 was dissolved in 1M methanesulfonic acid (MSA) (Sigma) then diluted in sterile ddH_2_O followed by adjusting pH value to 3-3.2 using 1M NaOH. Olaparib (100 mg/kg) or AZD9574 (3 mg/kg) was administered by oral gavage 1 h before radiation and continued for 6 days.

### Statistics

All data are presented as mean ± SEM unless otherwise stated. When assessing statistical significance between two treatment groups, continuous variables were analyzed using the unpaired Student *t-*test and Mann-Whitney test for normally and non-normally distributed data, respectively. In cases of more than two groups, ANOVA with the Tukey post-comparison test or Kruskal-Wallis analysis was used. Mouse survival analysis was examined using the log-rank test. P-values <0.05 were considered statistically significant and are denoted in the figures as follows: *P<0.05, **P<0.01, ***P<0.001, and ****P<0.0001. All tests were 2-sided. All statistical analyses were performed using GraphPad Prism 8 (GraphPad Inc.).

## Results

### H3K27M mutant cells are homologous recombination repair defective

The aberrant gene expression caused by H3K27M mutation may lead to dysfunctional DNA repair [48, 49]. The functional effect of H3K27M mutation on DNA repair, however, remains unclear. To identify targetable DNA repair vulnerabilities caused by H3K27M mutation, we used H3K27M isogenic DMG cell lines in which the spontaneous H3K27M mutation was edited by CRISPR/Cas9 to generate the knockout (KO) control (**Figure 1A**) [44]. Under both baseline conditions and in response to radiation, we observed more persistent γH2AX in H3K27M mutant cells relative to KO, supporting an H3K27M-mediated DNA repair deficiency (**Figure 1B, 1C, and Supplementary Figure S1A**). The delayed kinetics of this persistent DNA damage (16-24h) led us to hypothesize a defect in HRR which is a relatively slow DSB repair pathway. To investigate HRR activity, we examined RAD51, a recombinase required for HRR, and found that RAD51 focus formation was significantly reduced in H3K27M mutant (vs. KO) cells in response to RT, supporting an HRR defect (**Figure 1D**). Consistent with these results, we observed that U2OS cells harboring the H3K27M mutation exhibited a significant decrease in HRR efficiency relative to wild type (WT) histone H3 measured with a GFP-based reporter (DR-GFP) (**Figure 1E and Supplementary Figure S1B**). Furthermore, HRR reporter activity [50] was significantly reduced in H3K27M mutant BT245 cells compared to isogenic KO control cells (**Supplementary Figure S1C**). H3K27M mutant DMG cells were also more sensitive to the PARP inhibitors, further supporting a functional HRR defect caused by H3K27M mutation (**Supplementary Figure S1D**). Consistent with PARP inhibitor sensitivity [51], H3K27M mutant cells exhibited higher poly(ADP-ribose) (PAR) levels compared to KO cells that was not attributable to differences in PARP1 or PARG protein levels, the main enzymes for PAR synthesis and degradation, respectively (**Figure 1F**). Taken together, these results demonstrate that H3K27M mutation induces a BRCAness phenotype characterized by persistence of radiation-induced DNA damage, reduced RAD51 focus formation and HRR reporter activity, and increased sensitivity to PARP inhibitor. These findings support PARP as a therapeutically targetable genetic dependency in H3K27M DMG.

**Figure 1.**
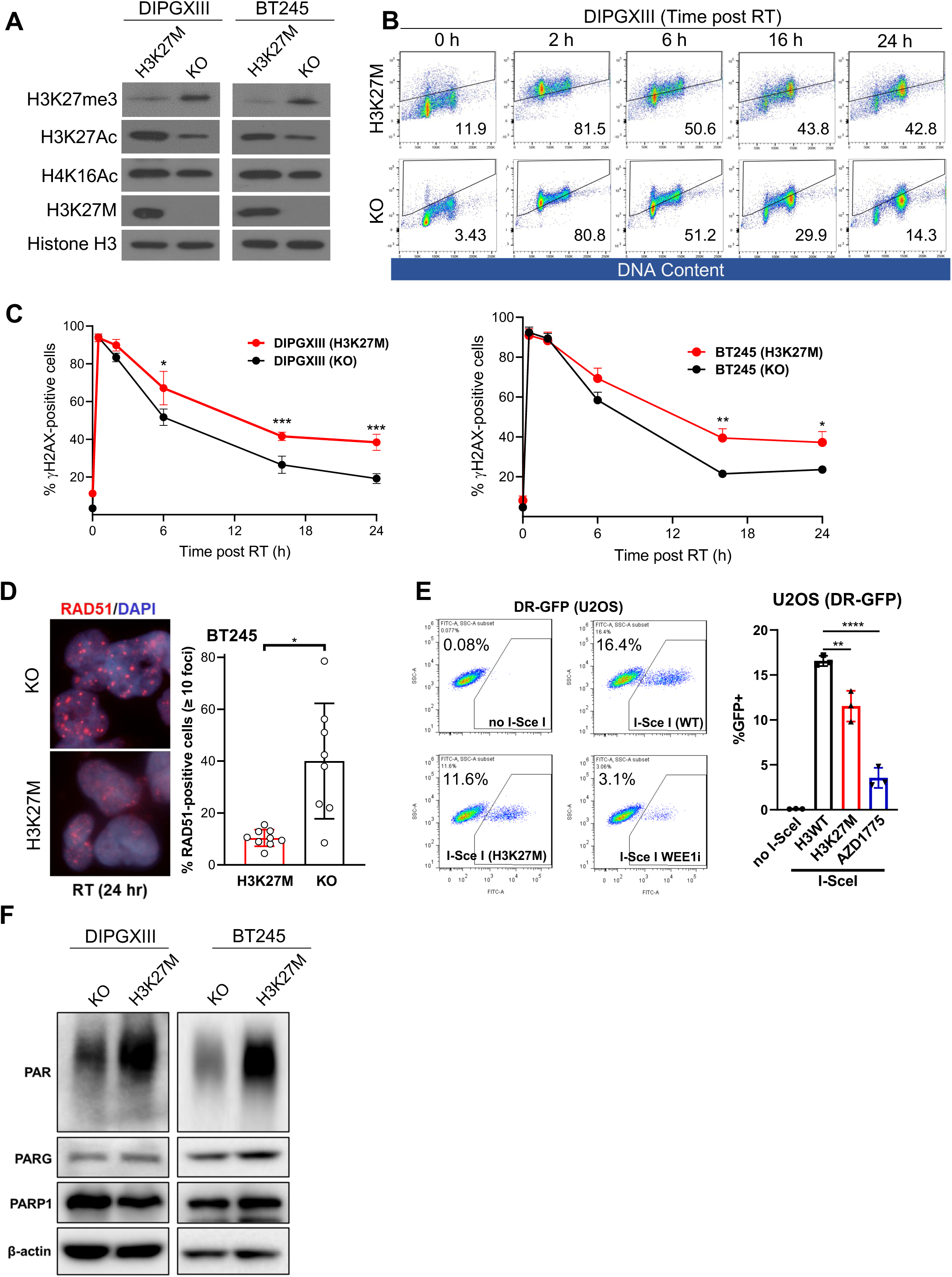
H3K27M causes a homologous recombination repair defect. (**A**) Western blot analysis of DIPGXIII and BT245 parental and their isogenic KO control cells with the indicated antibodies. Histone H3 represent loading controls. (**B**) H3K27M isogenic DIPGXIII cells were assessed for γH2AX positivity by flow cytometry at the indicated times post-RT (6 Gy). (**C**) Data plotted are the mean percentage of γH2AX positive cells ± SE (n = 3 independent experiments) in DIPGXIII (left) and BT245 (right) parental H3K27M and KO isogenic cells. (**D**) H3K27M isogenic BT245 cells were assessed for RAD51 foci by immunofluorescence at 24 h post-RT (6 Gy). Quantitative analysis data was plotted for RAD51 foci positive cells (n=8-10 fields). (**E**) DR-GFP U2OS cells with either H3K27M or WT histone H3 expression were infected with I-Sce I adenovirus. GFP expression (upon HR repair) was assessed by flow cytometry. Experiments was performed three times (±SEM). WEE1 inhibitor AZD1775 (100 nM) was used as positive control. (**F**) Western blot analysis of DIPGXIII and BT245 parental and their isogenic KO control cells with PAR, PARP1, and PARG antibodies. β-actin represent loading controls.

### H3K27M mutation abrogates RT-induced K63-linked polyubiquitination on histone H1 and HRR protein recruitment

To identify the underlying mechanism(s) of the HRR deficiency in H3K27M cells, we first assessed expression of genes involved in HRR genes by bulk RNA sequencing of H3K27M and isogenic KO BT245 and DIPGXIII cells [44] as H3K27M causes profound epigenetic dysregulation and gene expression alterations. Surprisingly, no significant difference in the expression levels of HRR genes was observed in the two isogenic paired cell lines (**Supplementary Figure S2A**). This is also supported by the analysis of PNOC (Pediatric Neuro-Oncology Consortium) database showing comparable expression of HRR genes between H3K27M DMG and WT pediatric high-grade glioma (pHGG) patients as well as no correlation of H3K27M mutation status with HRR (**Supplementary Figure S2B and S2C**) [52, 53], indicating H3K27M-mediated HRR deficiency is not caused by altered HRR gene expression.

Histone modification mediates the DNA damage responses (DDR) by facilitating the recruitment of DNA repair proteins to DNA damage sites. Gain-of-function mutation of H3K27M causes global loss of H3K27me2/3 marks with a *trans* enrichment of acetylation at multiple sites on core histones, including H3K27Ac and H4K16Ac [8, 9, 19] (**Figure 1A**). Histone acetylation may impede the linker histone H1 from binding to chromatin [21, 22], whereas DNA damage-dependent histone H1 non-proteolytic ubiquitination (via K63 linkage) generates binding sites for HRR proteins, such as BRCA1 and RAD51 [54]. As expected, we found decreased association of histone H1.2 with histone H3 in H3K27M DMG cells relative to isogenic KO cells (**Figure 2A**). In vitro protein binding assays using purified GST-H1.2 also showed reduced affinity with histone H3 in H3K27M mutant cells (**Figure 2B**). Further, H3K27M abrogated RT-induced accumulation of K63-linked ubiquitination of histone H1.2 in 293T cells (**Figure 2C**). In agreement, RT-induced K63 ubiquitylation was reduced in H3K27M DMG cells compared to KO control (**Figure 2D**). Upon DNA damage, E3 ubiquitin ligase RNF8, which is recruited by γH2AX/MDC1 axis, primes the formation of K63-linked polyubiquitination chains on H1, which subsequently relocates RNF168 via binding to its ubiquitin-interacting motifs (UIM) and leads to further propagation of K63-linked ubiquitination at DNA damage sites [55, 56]. To define the effect of H3K27M on the γH2AX/MDC1/RNF8/RNF168 axis, we assessed radiation-induced foci of MDC1, RNF8, and RNF168. While staining for MDC1 and RNF8 foci were comparable in H3K27M and KO cells, the relative amount of RNF168 was reduced in H3K27M cells (**Figure 2E**). To confirm this observation, we used a GFP fused reporter that can bind ubiquitinated H2AK13/15, which is modified by RNF168 (Reader1.0-eGFP) [57] and found a significant reduction in the recruitment of Reader1.0 to DNA damage sites in cells with H3K27M mutation (**Figure 2F**). Finally, RAP80 (receptor-associated protein 80) is a ubiquitin-binding protein that can specifically recognize K63-linked polyubiquitin chains, thus targeting the BRCA1 complex to DNA damage sites [58]. H3K27M DMG cells showed an impairment of RAP80/BRCA1/RAD51 loading to the chromatin (**Figure 2G**). Taken together, these data suggest that the HRR defect in H3K27M mutant cells is attributable to global hyper-acetylation of core histones consequently leading to H1 disassociation and subsequent loss of the K63 ubiquitination required for proper localization of HRR proteins to DSB sites (**Figure 2H**).

**Figure 2.**
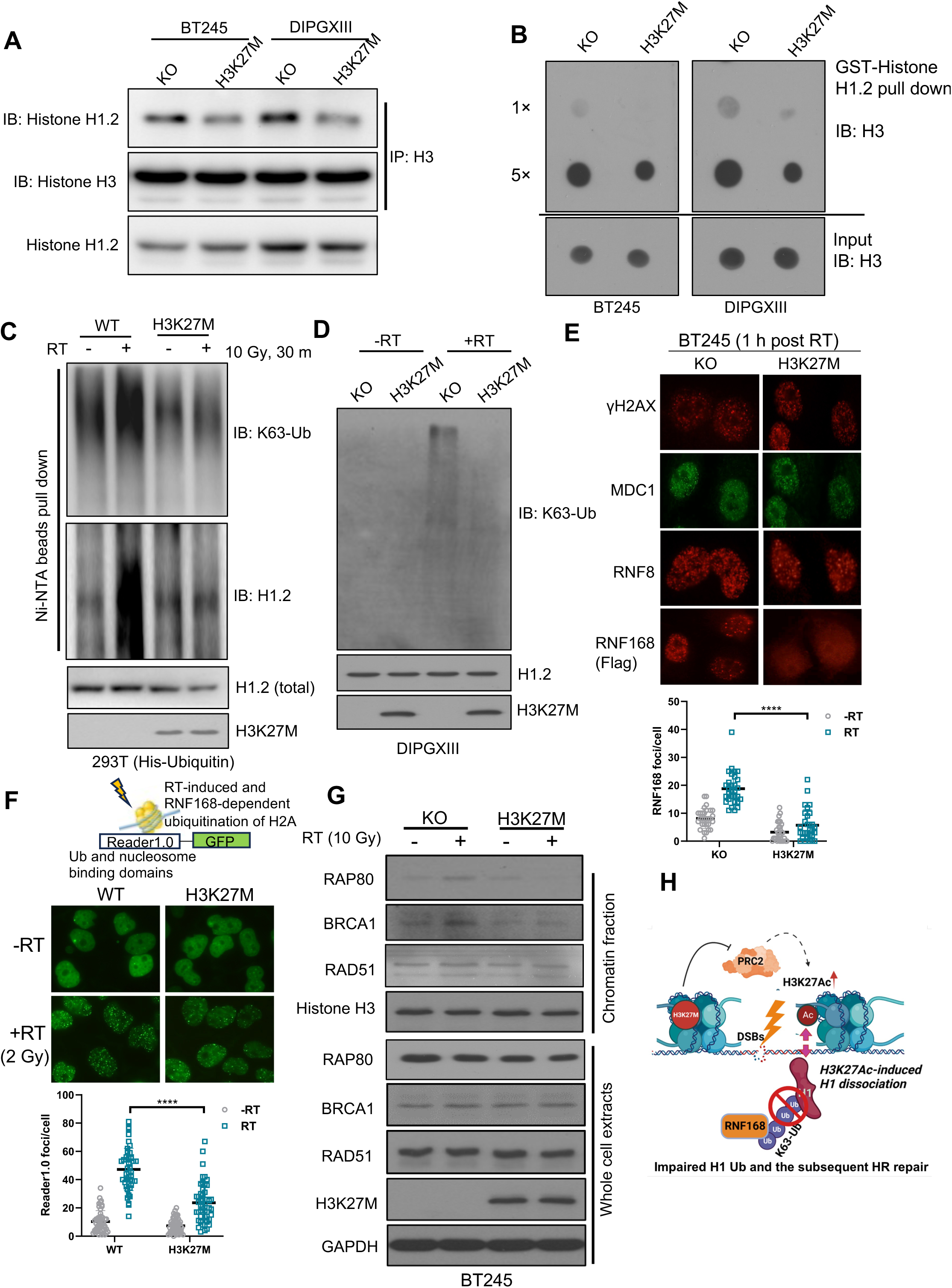
H3K27M mutation abrogates RT-induced K63-linked polyubiquitination of histone H1 and recruitment of HRR proteins. (**A**) Co-immunoprecipitation analysis of histone H3 and linker histone H1 interactions in BT245 and DIPGXIII parental and KO cells. (**B**) In vitro GST-pull down analysis of the binding of purified histone H1.2 with histone H3 extracted from H3K27M and isogenic KO BT245 and DIPGXIII cells. (**C**) H3K27M and WT 293T cells transfected with His-ubiquitin were harvested 30 min post-RT (10 Gy) for denatured Ni-NTA bead purification to isolate ubiquitinated proteins, followed by immunoblotting with H1.2 and K63 polyubiquitination antibodies. (**D**) Western blot analysis of DIPGXIII parental and their isogenic KO control cells with or without RT (10 Gy, 30 min) treatment with K63-Ub, H1.2 and H3K27M antibodies. (**E**) H3K27M and H3K27M KO BT245 cells (without or with FLAG-RNF168 transfection) were stained for γH2AX, MDC1, RNF8 or RNF168 (FLAG) at 1 hr post-RT (10 Gy) (upper). Data plotted are the number of FLAG-RNF168 foci per cell (n=30 cells). (**F**) H3K27M and WT Hela cells were infected with lentivirus expressing Reader1.0-GFP reporter were irradiated (2 Gy, 30 min) and monitored for reporter foci formation. (**G**) Western blotting analysis of RAP80, BRCA1, and RAD51 in the chromatin fractionation and total cell lysates in H3K27M and KO BT245 cells post-RT (10 Gy, 1h). (**H**) Schematic model showing that attenuated interaction of core histones and linker histone H1 in H3K27M cells leads to impairment of RT-induced K63 polyubiquitination and HRR protein recruitment.

### PARP inhibition selectively radiosensitizes H3K27M DMG cells, spheres, and tumors

PARP is vital for the repair of RT-induced single-strand DNA breaks (SSBs), the most abundant type of RT damage. PARP inhibition leads to the conversion of SSBs to cytotoxic DSBs in replicating cells which is synthetic lethal in HRR deficient cancer [59, 60]. Consistent with this concept, olaparib selectively sensitized human (BT245 and DIPGXIII) and mouse (PPK, p53R273H;PDGFRA;H3.3K27M) [46] H3K27M mutant DMGs to both low (2 Gy) and high (6 Gy) dose RT, but not isogenic KO or WT cells (**Figure 3A and Supplementary Figure S3A**). Radiosensitization by olaparib in BT245 and PPK cells was further supported in the 3D neurosphere formation assay (**Supplementary Figure S3B**). Olaparib and RT (6 Gy) caused more persistent DNA damage in H3K27M mutant DMG cells compared to matched KO cells, as evidenced by increased γH2AX and comet tail moment (**Figure 3B, 3C and Supplementary Figure S3C, S3D**). Consistent with the presence of unrepaired DSBs, BT245 and DIPGXIII cells arrested in G2/M following olaparib and radiation treatment (**Figure 3D and Supplementary Figure S3E**). Furthermore, olaparib and RT caused an increase in sub-G1 cells suggesting an apoptotic response that was confirmed by increased Annexin V/PI staining assay (**Figure 3E**) with no significant effect on senescence (**Supplementary Figure S3F**).

**Figure 3.**
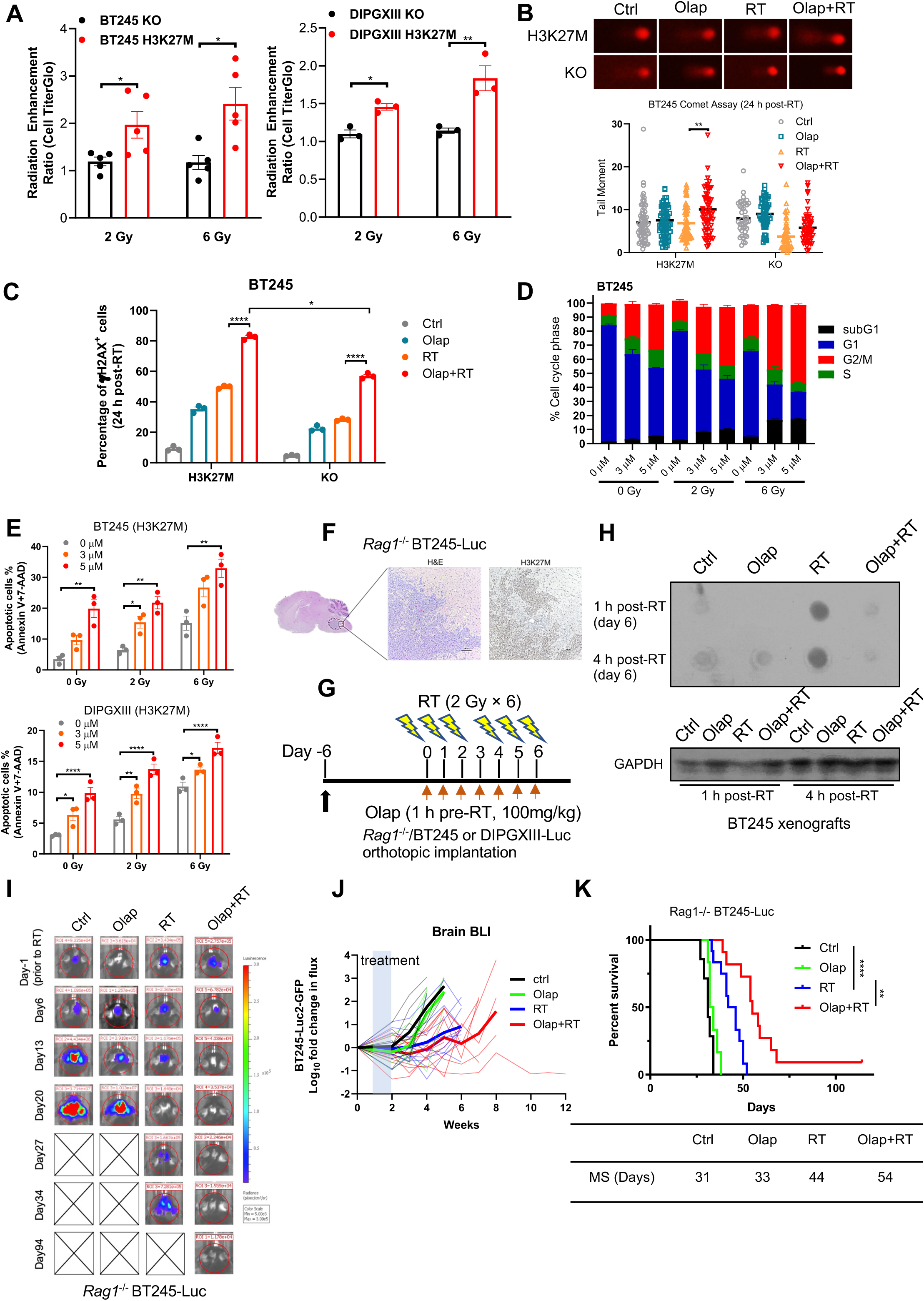
Selective radiosensitization of H3K27M DMG cells and orthotopic tumors by PARP inhibitor. (**A**) CellTiter Glo analysis of H3K27M and KO BT245 and DIPGXIII cell viability 5 days after olaparib (3 µM) and RT (2 Gy or 6 Gy) treatment. (**B**) Alkaline comet assay to assess DNA damage levels in H3K27M and isogenic KO BT245 cells in response to radiation (6 Gy) and/or olaparib (3 µM) treatment. (**C**) Data plotted are the mean percentage of γH2AX positive cells ± SEM (n = 3 independent experiments) in BT245 parental H3K27M and KO isogenic cells after 24 h of olaparib (3 µM) and RT (6 Gy) treatment. (**D**) Cell cycle distribution assessed by flow cytometry (propidium iodide) in BT245 cells with after treatment with radiation and olaparib as indicated. (**E**) Percentage of apoptotic cells assessed by Annexin V/7-AAD flow cytometry in BT245 and DIPGXIII cells after treatment with radiation and olaparib. (**F**) H&E-stained sections from the pons of *Rag1*^-/-^ mice implanted with BT245-Luciferase cells. Scale bars, 100 µm. (**G**) Schematic showing schedules of olaparib and radiation treatment. Olaparib (100 mg/kg) was orally administered approximately 1 hour before RT (2 Gy) on days 0–2 and 4–6. (**H**) Dot blotting analysis showing PAR levels in BT245 tumors that harvested either 1 h or 4 h after the last dose of RT. GAPDH represents loading control. (**I**) Representative in vivo bioluminescence images of BT245-Luciferase tumors taken before, during, and after treatment with olaparib and/or RT. (**J**) In vivo growth of BT245-Luciferase tumors of each animal (thin lines) and average (thick lines) bioluminescence intensities at each timepoint in response to olaparib and RT treatment. (**K**) Survival analysis of *Rag1*^-/-^ mice harboring BT245-Luciferase tumors treated as indicated (Ctrl n=7, Olap n=6, RT n=12, Olap+RT n=11). Data were analyzed using the log-rank test. MS, median survival.

To test the therapeutic efficacy of combined treatment with olaparib and RT in vivo, we orthotopically implanted BT245 or DIPGXIII cells with luciferase in the brainstem of *Rag1^-/-^* mice using the screw-guided method as illustrated (**Supplementary Figure S3G**) [47, 61]. Engrafted tumors within the pons were verified by histology and H3K27M immunohistochemistry (**Figure 3F**). We first assessed the ability of olaparib to inhibit PARP in orthotopic brain tumors. Treatment with olaparib reduced PAR levels following RT demonstrating that olaparib penetrates orthotopic DMG tumors and inhibits PARP (**Figure 3G and 3H**). To assess the therapeutic efficacy of olaparib and RT in H3K27M DMG, we monitored tumor burden by bioluminescence as well as overall survival in mice with BT245 or DIPGXIII tumors. Bioluminescent signals declined in response to RT or the combination of olaparib and RT in both the BT245 and DIPGXIII models (**Figure 3I and 3J, Supplementary Figure S3H and S3I**). Further, mice treated with the combination of olaparib and RT had longer overall survival compared to RT alone in both BT245 and DIPGXIII models with 10 and 7 day improvements in overall survival, respectively, and a 10% complete response rate (BT245 model) (**Figure 3K and Supplementary Figure S3J**). Treatment with olaparib and brain RT was well tolerated in terms of lack of significant weight loss and absence of neurotoxicity in normal brain tissues (**Supplementary Figure S3K and Supplementary Figure S3L**). Taken together, these studies demonstrate preferential radiosensitization of H3K27M DMG by PARP inhibition with a promising therapeutic index in vivo.

### PARP inhibition potentiates RT-induced type I interferon signaling in H3K27M DMG cells

Radiation stimulates the immunogenicity and visibility of tumor cells to cytotoxic immune cells, that can be further enhanced by combination with DDR inhibitors [28, 29]. To understand the effects of PARP inhibitor and RT on the immunogenicity of H3K27M mutant DMG in an unbiased manner, we performed bulk RNA-seq analysis and GSEA (Gene Set Enrichment Analysis). We found that many of the top enriched pathways were immune related including the innate immune response as the top enriched pathway in response to olaparib and RT (vs RT alone) (**Figure 4A**, **Table 1**). Owing to the key function of type I interferon (T1IFN) production and signaling in RT-induced antitumor immunity, we incorporated an *IFNβ1* promoter driven GFP reporter in H3K27M mutant DMG cells and their isogenic KO or KD controls (**Figure 4B**). While *IFNβ1*-GFP reporter expression was modestly increased by olaparib or RT treatment alone, combined treatment yielded a synergistic induction of *IFNβ1*-GFP that was maximal in the H3K27M mutant DMG cells versus control cells (**Figure 4B**). Consistent with increased reporter expression, olaparib in combination with RT also enhanced endogenous *IFNβ1* mRNA expression selectively in H3K27M DMG cells, noting a >100-fold induction of *IFNβ1* in H3K27M mutant DIPG007 cells in response to combined treatment (**Figure 4C and Supplementary Figure S4A**).

**Figure 4.**
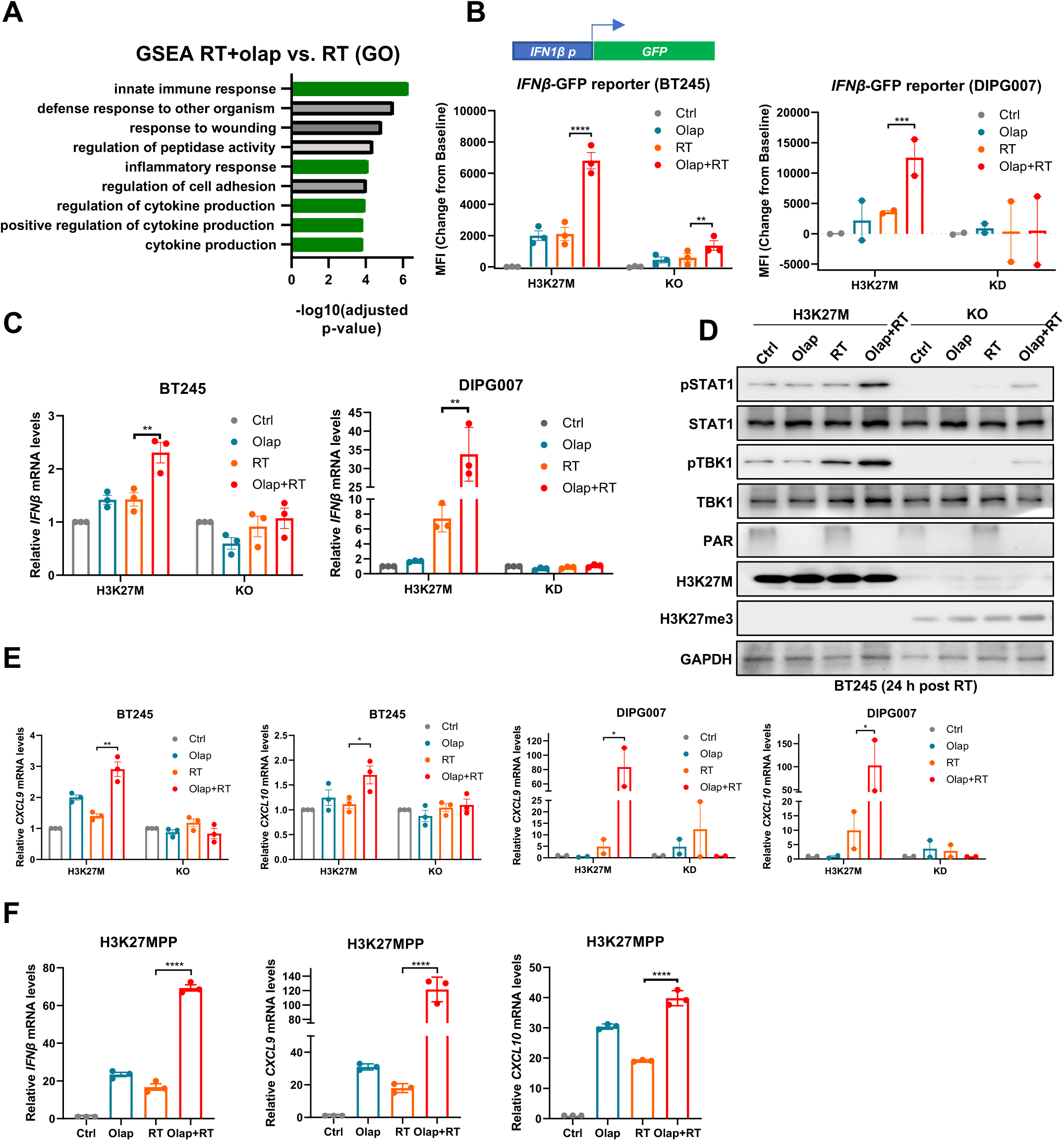
PARP inhibitor and RT preferentially induce type I interferon signaling in H3K27M DMG cells. (**A**) Gene ontology analysis run with clusterProfiler of differentiated genes in BT245 parental cells in response to RT and olaparib versus RT alone (72 h after treatment). (**B**) H3K27M and isogenic KO BT245 and KD DIPG007 cells with stable *IFNβ1* promoter-GFP reporter were treated as indicated and assessed for MFI of GFP expression. (**C**) qPCR for *IFNβ1* mRNA in H3K27M and KO BT245 and DIPG007 cells treated as indicated. (**D**) Western blot analysis of pTBK1, TBK1, pSTAT1, and STAT1 levels in H3K27M and KO BT245 cells at 24 hr after treated with olaparib and/or RT. (**E**) qPCR analysis of mRNA levels of interferon-stimulated genes *CXCL9* and *CXCL10* in BT245 and DIPG007 cells with or without H3K27M mutation at 72 h after olaparib and/or RT treatment. (**F**) qPCR analysis of *Ifnb*, *Cxcl9* and *Cxcl10* mRNA levels in mouse H3K27MPP DMG cells at 72 h following treatment with olaparib and/or RT.

Consistent with the increase in T1IFN, olaparib and RT activated T1IFN signaling marked by increased levels of phosphorylated TBK1 (S172) and STAT1 (Y701), upstream and downstream intermediates of T1IFN signaling, respectively, that were maximal in H3K27M mutant DMG cells (**Figure 4D and Supplementary Figure S4B**). In addition, expression of the interferon stimulated genes *CXCL9* and *CXCL10* was significantly increased by olaparib and RT in H3K27M mutant DMG cells (**Figure 4E and Supplementary Figure S4C**). Finally, we extended our studies to a murine H3K27M DMG model (H3K27MPP; generated from neuron progenitor cells infected with *H3K27M*, *p53^R273H^*, and *PDGFR* expression cassettes) and found similar to the human DMG models that olaparib and RT significantly increased expression *IFNβ1*, *CXCL9*, and *CXCL10* mRNA that was accompanied by modest induction of pTBK1 and pSTAT1 (**Figure 4F and Supplementary Figure S4D**) [45]. Taken together, these data show that PARP inhibition potentiates RT-induced immunogenicity through increased T1IFN expression, interferon stimulated gene response, and T1IFN signaling preferentially in H3K27M DMG cells.

### PARP inhibitor and RT induce NKG2D activating ligand expression and NK cell-mediated lysis

T1IFN is necessary for the proper expansion, maturation, and activation of both T and NK cells [62, 63]. Although cytotoxic T cells play a central role in antitumor immune responses, NK cells have the potential for greater cytotoxic activity in DMG and correlate with overall survival in DMG patients [34, 37]. A key feature of NK cells is their ability to recognize stressed “self” cells through the activation of NK cell surface receptors like NKG2D that when engaged by respective ligands (e.g., MICA, MICB, and ULBP1-6 in human; Rae, H60, and Mult1 in mouse) induce NK cell degranulation, cytokine production, and NK cell-mediated lysis [40]. To evaluate the role of NK cells in the therapeutic efficacy of olaparib and RT, we first assessed the NKG2D activation ligands MICA/B and ULBP-3 which are expressed in DMG tumors [34]. Treatment with olaparib and RT caused a 5-10-fold increase of MICA/B and ULBP-3 cell surface expression in H3K27M DMG cells (**Figure 5A**) that was also accompanied by increased mRNA expression of *MICA, MICB,* and *ULBP-3* mRNAs (**Figure 5B and Supplementary** Figure 5A). Furthermore, expression of the analogous murine NKG2D ligands, Rae1 and H60, was increased by olaparib and RT in the murine PPK H3K27M mutant DMG model (**Figure 5C**). This effect was specific to PARP inhibition, as NKG2D ligand expression was not enhanced by inhibition of other DDR proteins such as ATM and ATR in combination with RT radiation (**Supplementary Figure S5B**). Instead, and consistent with a prior study [41, 64], the induction of NKG2D ligands by olaparib and RT was dependent on ATM and ATR activity, as simultaneous inhibition of both ATM and ATR prevented the induction of MICA/B in BT245 and DIPG007 cells (**Figure 5D**). In contrast, NKG2D ligand expression in response to combined treatment was not dependent on T1IFN signaling as neither STING inhibitor (H151) nor T1IFN receptor blocking antibody reduced NKG2D ligand induction (**Supplementary Figure S5C**). Because NK cell activity is also regulated by inhibitory receptors, we assessed the effects of olaparib and RT on the expression of two inhibitory ligands HLA-E and HLA-A/B/C which bind NKG2A/p94 and KIRs, respectively, to repress NK cell activity. We found no induction of these inhibitory ligands by olaparib and RT (**Supplementary Figure S5D**). These results indicate a shift in the repertoire of NK regulatory proteins on the surface of H3K27M mutant DMG cells in favor of stimulation of cytotoxic NK cells.

**Figure 5.**
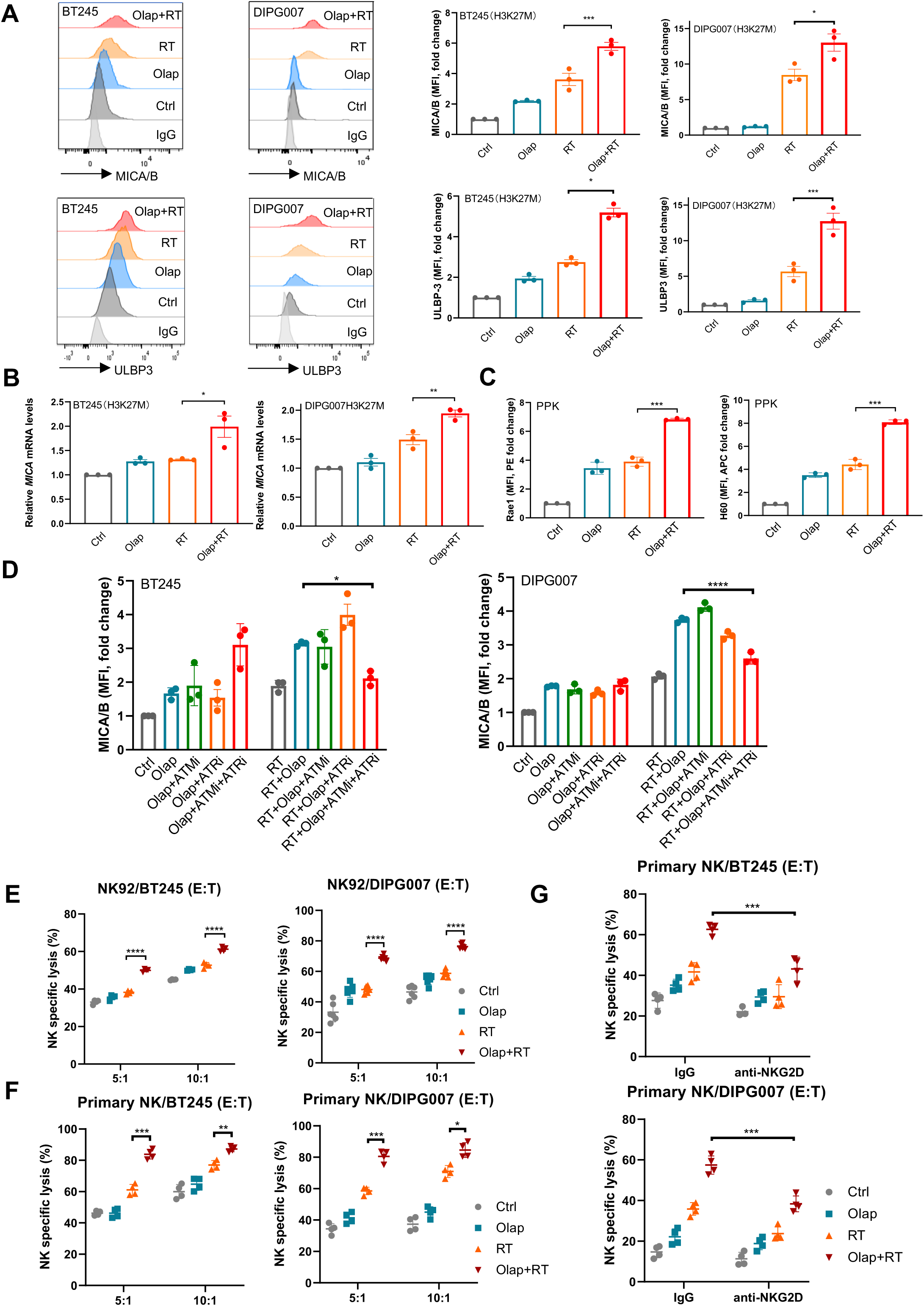
PARP inhibitor and RT increase expression of NKG2D activating ligands and induce NK cell lysis. (**A**) Flow cytometry analysis of cell surface MICA/B and ULBP-3 protein levels in BT245 and DIPG007 cells at 72 h following indicated treatment. (**B**) qPCR for *MICA* mRNA levels in BT245 and DIPG007 cells at 72 h after olaparib and/or RT treatment. (**C**) Flow cytometry analysis of cell surface Rae1 and H60 levels in H3K27M mouse DMG cells at 72 h after olaparib and/or RT treatment. (**D**) Cell surface MICA/B levels of H3K27M parental BT245 and DIPG007 cells in response to RT and/or DDR inhibitor treatment as indicated. (**E, F**) Lysis of target BT245 and DIPG007 cells co-cultured with NK92 (E) or primary activated NK cells (F) after 72 h pre-treatment of DMG cells with olaparib (3 µM) and/or RT (8 Gy). (**G**) Effect of NKG2D blocking antibodies on NK cell killing of BT245 and DIPG007 cells with olaparib (3 µM) and/or RT (8 Gy) treatment.

To assess whether olaparib and RT enhance NK cell-mediated lysis of H3K27M mutant DMG cells, BT245 and DIPG007 cells treated with olaparib and RT were fluorescently labeled with calcein-AM dye and then co-cultured with human NK-92 effector cells, an FDA-approved NK immunotherapy cell line derived from NK cell lymphoma, at two different effector to target (E:T) ratios (5:1 and 10:1). Combination treatment of DMG cells with olaparib and RT led to increased lysis of both DMG cells lines (**Figure 5E**). Further, using primary NK cells (**Supplementary Figure S5E**), we found similar enhancement of NK cell-mediated killing of DMG cell lines treated with PARP inhibitor and RT (**Figure 5F**). As anticipated, NK-cell mediated lysis of DMG cells was reversed by the addition of an NKG2D blocking antibody further supporting the necessity of NKG2D activation for NK cell-mediated lysis (**Figure 5G**). To address the selectivity of NK cell-mediated lysis of DMG tumor cells, we conducted parallel studies in normal human astrocytes (NHA). PARP inhibitor did not enhance MICA/B expression or NK cell-mediated lysis of DMG cells following RT (**Supplementary Figure S5F and S5G**). Collectively, these results demonstrate that treatment with PARP inhibitor and RT stimulates NK activating ligand expression in H3K27M DMG cells leading to selective lysis of DMG tumor cells by activated NK cells.

### NK cells are required for the therapeutic efficacy of PARP inhibitor and RT in H3K27M DMG

To investigate the contribution of the immune system and specifically NK cells to therapeutic responses in H3K27M DMG, we established an immunocompetent syngeneic orthotopic DMG model consisting of H3K27MPP cells implanted into the brainstem of C57BL/6 mice (**Figure 6A**). To directly assess the contribution of NK cells to the therapeutic efficacy of olaparib and radiation, mice were simultaneously treated with anti-NK1.1 to deplete NK cells or with control anti-IgG as illustrated (**Figure 6A**). We found in mice with NK cells intact, that combined treatment with olaparib and RT produced a median survival of 41 days. Importantly, in mice depleted of NK cells, the median survival following olaparib and RT treatment was significantly reduced (22 days) indicating the requirement for NK cells in the therapeutic efficacy of PARP inhibitor and RT (**Figure 6B**).

**Figure 6.**
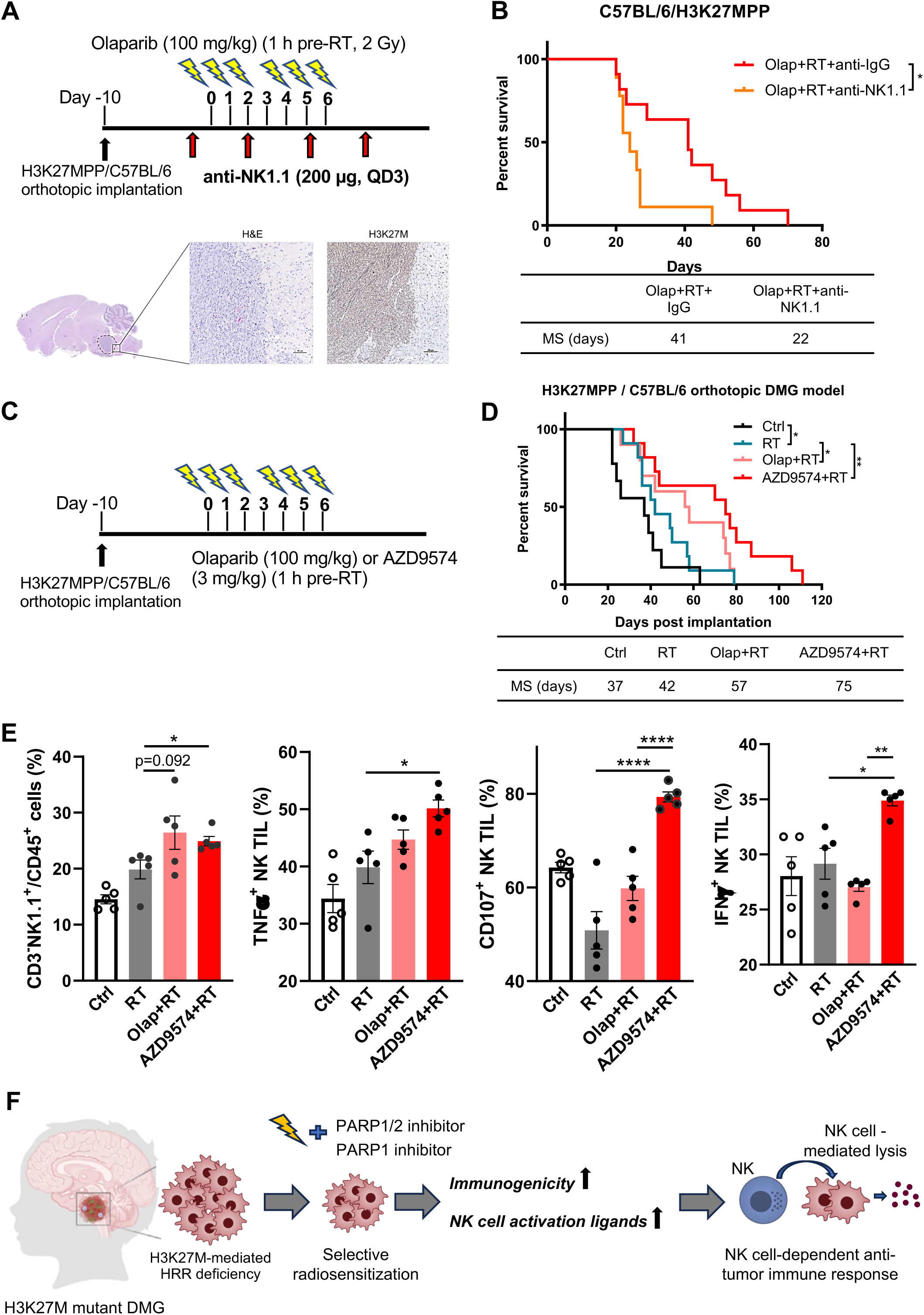
PARP inhibitors and RT induce a therapeutic response in immune competent mice that is dependent upon NK cells. (**A**) Schematic showing schedule of olaparib, RT, and anti-NK1.1 antibody treatment. Olaparib (100 mg/kg) was orally administered approximately 1 hour before each fractionated dose of RT (2 Gy, 6 fractions) on days 0–2 and 4–6. Mouse anti-NK1.1 antibody (200 μg/mL) was intraperitoneally injected every 3 days for a total of 4 doses (upper). H&E-stained sections from C57BL/6 mice with H3K27M DMG tumors. Scale bars, 100 µm (lower). (**B**) Survival analysis of C57BL/6/H3K27MPP DMG mice treated as indicated in 6A (RT+olap+IgG n=11, RT+olap+anti-NK1.1 n=9). Data were analyzed using the log-rank test. (**C**) Schematic showing schedules of olaparib or AZD9574 and radiation treatment. Olaparib (100 mg/kg) and AZD9574 (3 mg/kg) was orally administered approximately 1 hour before RT (2 Gy) on days 0–2 and 4–6. (**D**) Survival analysis of C57BL/6/H3K27MPP DMG mice treated as indicated (Ctrl n=9, RT n=11, RT+olap n=10, RT+AZD9574 n=11). Data were analyzed using the log-rank test. (**E**) Flow cytometry analysis of intratumoral NK cells (in the CD45^+^ cell populations), and TNF-α^+^, CD107^+^, and IFNγ^+^ NK cells from mouse H3K27MPP DMG tumors treated as indicated. Data are the mean ± SEM (n = 5 mice/group). (**F**) Proposed model of H3K27M mutation-mediated HRR deficiency that leads to selective radiosensitization by PARP inhibition and the unique characters of RT and PARP inhibitors in NK cell activation in DMG.

Beyond establishing the importance of NK cells to therapeutic efficacy, we next sought to optimize our therapeutic strategy by incorporating the novel blood-brain barrier penetrant PARP1 selective inhibitor, AZD9574 [43]. Mice with orthotopic H3K27MPP tumors were treated with RT alone or in combination with olaparib or AZD9574 as illustrated (**Figure 6C**). In the absence of treatment, the median survival in this model was 37 days. Radiotherapy alone extended the median survival to 42 days (**Figure 6D**). The addition of olaparib to RT further extended median survival to 57 days which was statistically significant in comparison to RT alone. To further optimize therapeutic efficacy, we also tested AZD9574 in combination with RT and found maximum therapeutic efficacy with a median survival reaching 75 days, that was tolerable as evidenced by a lack of weight loss and no appreciable normal brain toxicity (**Figure 6D and Supplementary Figure S6A and S6B**). We then investigated the effects of olaparib or AZD9574 with RT on intratumoral cytotoxic immune cells, including both CD8^+^ T cells as well as NK cells the latter of which were detectable at baseline in orthotopic DMG tumors (**Supplementary Figure S6C**). Combined treatment with PARP inhibitor and RT did not consistently increase CD8^+^ T cell frequency or activation status (**Supplementary Figure S6D and S6E**). In contrast, PARP inhibitor significantly increased intratumoral NK cell frequency (CD3^-^NK1.1^+^CD45^+^) and activity (TNFα^+^, CD107^+^, or IFNγ^+^) (**Figure 6E and Supplementary Figure S6F**). Moreover, AZD9574 had a stronger effect than olaparib in activating intratumoral NK cells following RT which may be a reflection of improved brain penetrance and/or PARP1 selectivity (**Figure 6E**). Collectively, these data support a model (**Figure 6F**) in which PARP inhibitor and RT promote NK cell cytotoxic functions in the TME of H3K27M DMG leading to therapeutic responses with the PARP1 selective inhibitor AZD9574 comparing favorably to PARP1/2 inhibition by olaparib.

## Discussion

H3K27M mutant histone is an oncogene in DMG but not a direct therapeutic target. This mutation disrupts the normal function of core histones and consequently is implicated in altered DNA repair [48, 49]. Here we show that H3K27M causes HRR deficiency by the “hyperacetylated” core histone-induced linker histone H1 dissociation leading to impairment of K63 polyubiquitination of H1 and RNF168/BRCA1/RAD51 protein recruitment, consistent with the key role of chromatin structure and histone modification in HRR. Our study directly connects a functional interaction between core histones and linker histone H1 in H3K27M mutant DMG. In addition to RAP80/BRCA1 recruitment, 53BP1 relocation is dependent on RNF168-catalyzed ubiquitination of histones H2A/H2AX on K13/15. The failure of RNF168 foci formation and H2A/H2AX (K13/K15) ubiquitination (Read1.0-GFP) in H3K27M mutant cells is also consistent with the previous study showing that H3K27M mutation caused an impairment of 53BP1 recruitment, supporting a profound effect of H3K27M on DNA repair [65].

Although H3K27M mutant DMG is defective in DSB repair, H3K27M DMG patient tumors are not especially sensitive to RT with outcomes characterized by an initial response followed by recurrence with poorer survival compared to patients with H3K27 wildtype high grade glioma [52, 66]. This study supports that H3K27M-mediated HRR deficiency is an actionable vulnerability that increases radiosensitization by PARP inhibitor. H3K27M mutation also increases the sensitivity of DMG cells to PARP inhibitor monotherapy (Figure S1D), an effect that we did not recapitulate in vivo likely due to the relatively ‘brief’ radiosensitization treatment schedule in comparison to the prolonged administration schedules required for PARP inhibitor monotherapy efficacy in HRR defective cancer [67]. Beyond the interaction of PARP and RT in HRR defective DMG, PARP1 expression is increased in DMG patients compared to normal tissues [68, 69]. The higher poly(ADP-ribosyl) levels in H3K27M versus KO cells further highlights that PARP is a promising target in H3K27M DMG. Clinically, however, the PARP1/2 inhibitor veliparib in combination with RT did not show a survival benefit in DMG patients [70]. A significant portion of patients on this trial did not have a decrease in PARP activity measured by PAR in PBMCs in response to veliparib nor did PAR reduction correlate with patient survival. It is unclear why this trial failed to produce clinical benefit, but poor brain penetrance or less potent PARP trapping (discussed below) may have contributed.

Here we show that the blood-brain barrier penetrant and potent PARP1 selective inhibitor AZD9574 [43] compares favorably to the first-generation PARP1/2 inhibitor olaparib to activate NK cells and prolong survival in a syngeneic orthotopic model. There are several potential contributing factors to the difference in efficacy of PARP1 versus PARP1/2 inhibition. PARP1 is the major mediator of cellular PARylation accounting for the majority of cellular PARylation in response to DNA damage [71]. In addition, PARP1 retention at DNA damage sites known as *PARP trapping* occurs in response to PARP inhibitors like olaparib and AZD9574 and is a key mechanism of radiosensitization (as well as monotherapy efficacy) of PARP inhibitors [43, 72]. In contrast, PARP2 may not be required for anticancer activity [73]. PARP2 has been shown to play a key role in the survival of hematopoietic cells and in the maintenance of cytotoxic T cell and NK cell activity [74, 75]. Future studies will define the relative contributions of these mechanisms to the therapeutic efficacy of PARP inhibition with radiotherapy with a focus on the immunomodulatory effects in the TME of H3K27M mutant DMG through global analyses (eg, single cell RNA-seq and spatial molecular imaging analysis).

We and others have shown that inhibition of the DDR can further enhance RT-induced tumor immunogenicity via immunogenic DNA damage that activates intrinsic nucleic acid sensors [25, 28, 30]. Here, we found a preferential induction of T1IFN expression and signaling in response to PARP inhibition and RT in the parental H3K27M mutant DMG cells relative to their isogenic KO controls, which may be attributable to the open chromatin structure and immunogenic DNA damage in H3K27M mutant cells. T1IFNs execute antitumor functions mainly through stimulating immune cells, especially CD8^+^ T cells and NK cells. Although DMG tumor cells express major histocompatibility complex class I (MHC-I), NK cells have the potential for greater cytotoxic activity in DMGs compared to cytotoxic T cells [34]. RT and other genotoxic agents are potent stimulators of NK cell activating receptor NKG2D ligand expression on tumor cells [41, 42]. Moreover, this induction is dependent on the ATM/ATR/CHK1/2 pathway, which makes PARP inhibitors unique in potentiating RT-induced NKG2D ligand expression and NK cell lysis. We found that antibody-mediated NKG2D blockade abrogated NK cell killing of DMG cells following PARP inhibitor and RT, indicating the critical role of NKG2D ligand induction in NK cell lysis efficacy. While NKG2D ligand expression following PARP inhibitor and RT did not require T1IFN signaling, it is still likely that these signaling axes cooperate to mediate the antitumor immune response.

The tumor microenvironment of syngeneic orthotopic H3K27M DMG tumors does not fully recapitulate that of human DMG, which is characterized by an immune naïve tumor microenvironment with a paucity of lymphocytes. Using a genetically engineered H3K27M DMG model could better provide a more clinically relevant model for understanding the effects of PARP1 and PARP1/2 inhibitors with radiotherapy on the TME [76]. On the other hand, NK directed immunotherapy or NK cellular therapy (eg, CAR-NK) offer alternative strategies for overcoming the immune naïve TME of DMG that should be further enhanced by selective *priming* of H3K27M mutant DMG tumor cells with PARP inhibitor and RT.

In conclusion, our study elucidates a novel connection between H3K27M mutation and HRR deficiency leading to selective radiosensitization of H3K27M mutant DMG by PARP inhibition (**Figure 6F**). Radiosensitization is associated with an enhanced innate immune response and increased expression of NK activating ligands in DMG cells. Consistent with increased immunogenicity, PARP inhibitor and RT promote selective NK cell lysis of DMG tumors cells (but not normal astrocytes), therapeutic responses in immune competent DMG tumor models, and increased NK cell infiltration and activation that is required for therapeutic efficacy. Finally, these studies support the potential for PARP1 selective inhibitors to provide improved efficacy over PARP1/2 inhibitors in inducing an NK cell-mediated antitumor immune response and support future clinical development of PARP inhibitor with RT in H3K27M mutant DMG, with the possibility for combination with NK cellular therapy or immunotherapy.

## Authorship

Conception and design of the study: M.A.M., Q.Z. Experimental design: Y. G., Z. L., L.A.P., M.A.M., Q.Z. Cell assay and molecular experiments: Y.G., Z.L., L.A.P., Z.W., J.D.P., A.D.

Bioinformatics: S.T., N.H., V.M.V. Mouse studies: Y.G., Z.L., Z.W., R.D., E.P. Acquisition, analysis, and/or interpretation of data: All authors. Writing original draft preparation: Y.G., M.A.M., Q.Z. Writing-review and editing: All authors. Study supervision: T.S.L., C.K., M.A.M., Q.Z. All authors discussed and reviewed the manuscript and approved the manuscript for publication.

## Data Availability

RNAseq data have been deposited in GEO (GSE268375). All data files are available from the corresponding authors upon request.

## Acknowledgements

We are grateful to Dr. Nada Jabado for providing BT245 and DIPGXIII parental and H3K27M KO isogenic cells. We thank Dr. Efrat Shema for the pInducer20/H3.3 WT or H3.3-K27M constructs, Drs. Tingting Yao and Robert Cohen for the Reader1.0-GFP reporter; Drs. Heng Lin and Amanda Huber for the flow cytometry analysis and human NK cell isolation.

## References

1. Cooney, T., et al., Contemporary survival endpoints: an International Diffuse Intrinsic Pontine Glioma Registry study. Neuro Oncol, 2017. 19(9): p. 1279–1280.

2. Louis, D.N., et al., The 2021 WHO Classification of Tumors of the Central Nervous System: a summary. Neuro Oncol, 2021. 23(8): p. 1231–1251.

3. Cohen, K.J., N. Jabado, and J. Grill, Diffuse intrinsic pontine gliomas-current management and new biologic insights. Is there a glimmer of hope? Neuro Oncol, 2017. 19(8): p. 1025–1034.

4. Hargrave, D., U. Bartels, and E. Bouffet, Diffuse brainstem glioma in children: critical review of clinical trials. Lancet Oncol, 2006. 7(3): p. 241–8.

5. Freeman, C.R., et al., Final results of a study of escalating doses of hyperfractionated radiotherapy in brain stem tumors in children: a Pediatric Oncology Group study. Int J Radiat Oncol Biol Phys, 1993. 27(2): p. 197–206.

6. Janssens, G.O., et al., Survival benefit for patients with diffuse intrinsic pontine glioma (DIPG) undergoing re-irradiation at first progression: A matched-cohort analysis on behalf of the SIOP-E-HGG/DIPG working group. Eur J Cancer, 2017. 73: p. 38–47.

7. Tinkle, C.L., et al., Radiation dose response of neurologic symptoms during conformal radiotherapy for diffuse intrinsic pontine glioma. J Neurooncol, 2020. 147(1): p. 195–203.

8. Schwartzentruber, J., et al., Driver mutations in histone H3.3 and chromatin remodelling genes in paediatric glioblastoma. Nature, 2012. 482(7384): p. 226–31.

9. Wu, G., et al., Somatic histone H3 alterations in pediatric diffuse intrinsic pontine gliomas and non-brainstem glioblastomas. Nat Genet, 2012. 44(3): p. 251–3.

10. Justin, N., et al., Structural basis of oncogenic histone H3K27M inhibition of human polycomb repressive complex 2. Nat Commun, 2016. 7: p. 11316.

11. Lewis, P.W., et al., Inhibition of PRC2 activity by a gain-of-function H3 mutation found in pediatric glioblastoma. Science, 2013. 340(6134): p. 857–61.

12. Mavragani, I.V., et al., Ionizing Radiation and Complex DNA Damage: From Prediction to Detection Challenges and Biological Significance. Cancers (Basel), 2019. 11(11).

13. Mendez-Acuna, L., et al., Histone post-translational modifications in DNA damage response. Cytogenet Genome Res, 2010. 128(1-3): p. 28–36.

14. Siddaway, R., et al., Oncohistone interactome profiling uncovers contrasting oncogenic mechanisms and identifies potential therapeutic targets in high grade glioma. Acta Neuropathol, 2022. 144(5): p. 1027–1048.

15. Katagi, H., et al., Radiosensitization by Histone H3 Demethylase Inhibition in Diffuse Intrinsic Pontine Glioma. Clin Cancer Res, 2019. 25(18): p. 5572–5583.

16. Watanabe, J., et al., Brd4 Inhibition as a Radiosensitizer through Blocking DNA Repair for the Treatment of Diffuse Midline Glioma. Neuro-Oncology, 2023. 25.

17. Watanabe, J., et al., BET bromodomain inhibition potentiates radiosensitivity in models of H3K27-altered diffuse midline glioma. J Clin Invest, 2024.

18. An, S., et al., Histone tail analysis reveals H3K36me2 and H4K16ac as epigenetic signatures of diffuse intrinsic pontine glioma. J Exp Clin Cancer Res, 2020. 39(1): p. 261.

19. Harpaz, N., et al., Single-cell epigenetic analysis reveals principles of chromatin states in H3.3-K27M gliomas. Mol Cell, 2022. 82(14): p. 2696–2713 e9.

20. Perry, C.A. and A.T. Annunziato, Influence of histone acetylation on the solubility, H1 content and DNase I sensitivity of newly assembled chromatin. Nucleic Acids Res, 1989. 17(11): p. 4275–91.

21. Perry, C.A. and A.T. Annunziato, Histone acetylation reduces H1-mediated nucleosome interactions during chromatin assembly. Exp Cell Res, 1991. 196(2): p. 337–45.

22. Ridsdale, J.A., et al., Histone acetylation alters the capacity of the H1 histones to condense transcriptionally active/competent chromatin. J Biol Chem, 1990. 265(9): p. 5150–6.

23. Deng, L., et al., STING-Dependent Cytosolic DNA Sensing Promotes Radiation-Induced Type I Interferon-Dependent Antitumor Immunity in Immunogenic Tumors. Immunity, 2014. 41(5): p. 843–52.

24. Falcke, S.E., et al., Clinically Relevant Radiation Exposure Differentially Impacts Forms of Cell Death in Human Cells of the Innate and Adaptive Immune System. Int J Mol Sci, 2018. 19(11).

25. Harding, S.M., et al., Mitotic progression following DNA damage enables pattern recognition within micronuclei. Nature, 2017. 548(7668): p. 466–470.

26. Valvo, V., et al., Editorial: Targeting DNA damage response to enhance antitumor innate immunity in radiotherapy. Front Oncol, 2023. 13: p. 1257622.

27. Haase, S., et al., H3.3-G34 mutations impair DNA repair and promote cGAS/STING-mediated immune responses in pediatric high-grade glioma models. J Clin Invest, 2022. 132(22).

28. Wang, W., et al., DNA-PK Inhibition and Radiation Promote Antitumoral Immunity through RNA Polymerase III in Pancreatic Cancer. Mol Cancer Res, 2022. 20(7): p. 1137–1150.

29. Zhang, Q., et al., Inhibition of ATM Increases Interferon Signaling and Sensitizes Pancreatic Cancer to Immune Checkpoint Blockade Therapy. Cancer Res, 2019. 79(15): p. 3940–3951.

30. Zhang, Q., et al., Potentiating the radiation-induced type I interferon antitumoral immune response by ATM inhibition in pancreatic cancer. JCI Insight, 2024. 9(6).

31. Baude, J., et al., Combining radiotherapy and NK cell-based therapies: The time has come. Int Rev Cell Mol Biol, 2023. 378: p. 31–60.

32. Gough, M.J. and M.R. Crittenden, The paradox of radiation and T cells in tumors. Neoplasia, 2022. 31: p. 100808.

33. Patin, E.C., et al., Harnessing radiotherapy-induced NK-cell activity by combining DNA damage-response inhibition and immune checkpoint blockade. J Immunother Cancer, 2022. 10(3).

34. Lieberman, N.A.P., et al., Characterization of the immune microenvironment of diffuse intrinsic pontine glioma: implications for development of immunotherapy. Neuro Oncol, 2019. 21(1): p. 83–94.

35. Lin, G.L., et al., Non-inflammatory tumor microenvironment of diffuse intrinsic pontine glioma. Acta Neuropathol Commun, 2018. 6(1): p. 51.

36. Persson, M.L., et al., The intrinsic and microenvironmental features of diffuse midline glioma: Implications for the development of effective immunotherapeutic treatment strategies. Neuro Oncol, 2022. 24(9): p. 1408–1422.

37. Bailey, C.P., et al., Pharmacologic inhibition of lysine-specific demethylase 1 as a therapeutic and immune-sensitization strategy in pediatric high-grade glioma. Neuro Oncol, 2020. 22(9): p. 1302–1314.

38. Vivier, E., et al., Natural killer cell therapies. Nature, 2024. 626(8000): p. 727–736.

39. Wolf, N.K., D.U. Kissiov, and D.H. Raulet, Roles of natural killer cells in immunity to cancer, and applications to immunotherapy. Nat Rev Immunol, 2023. 23(2): p. 90–105.

40. Raulet, D.H., et al., Regulation of ligands for the NKG2D activating receptor. Annu Rev Immunol, 2013. 31: p. 413–41.

41. Gasser, S., et al., The DNA damage pathway regulates innate immune system ligands of the NKG2D receptor. Nature, 2005. 436(7054): p. 1186–90.

42. Weiss, T., et al., NKG2D-Dependent Antitumor Effects of Chemotherapy and Radiotherapy against Glioblastoma. Clin Cancer Res, 2018. 24(4): p. 882–895.

43. Staniszewska, A.D., et al., Preclinical Characterization of AZD9574, a Blood-Brain Barrier Penetrant Inhibitor of PARP1. Clin Cancer Res, 2024. 30(7): p. 1338–1351.

44. Harutyunyan, A.S., et al., H3K27M induces defective chromatin spread of PRC2-mediated repressive H3K27me2/me3 and is essential for glioma tumorigenesis. Nat Commun, 2019. 10(1): p. 1262.

45. Golbourn, B.J., et al., Loss of MAT2A compromises methionine metabolism and represents a vulnerability in H3K27M mutant glioma by modulating the epigenome. Nat Cancer, 2022. 3(5): p. 629–648.

46. Miklja, Z., et al., Everolimus improves the efficacy of dasatinib in PDGFRalpha-driven glioma. J Clin Invest, 2020. 130(10): p. 5313–5325.

47. Marigil, M., et al., Development of a DIPG Orthotopic Model in Mice Using an Implantable Guide-Screw System. PLoS One, 2017. 12(1): p. e0170501.

48. Pedersen, H., K. Schmiegelow, and P. Hamerlik, Radio-Resistance and DNA Repair in Pediatric Diffuse Midline Gliomas. Cancers (Basel), 2020. 12(10).

49. Parsels, L.A., et al., Developing H3K27M mutant selective radiosensitization strategies in diffuse intrinsic pontine glioma. Neoplasia, 2023. 37: p. 100881.

50. Arnoult, N., et al., Regulation of DNA repair pathway choice in S and G2 phases by the NHEJ inhibitor CYREN. Nature, 2017. 549(7673): p. 548–552.

51. Bianchi, A., et al., PARP-1 activity (PAR) determines the sensitivity of cervical cancer to olaparib. Gynecol Oncol, 2019. 155(1): p. 144–150.

52. Mackay, A., et al., Integrated Molecular Meta-Analysis of 1,000 Pediatric High-Grade and Diffuse Intrinsic Pontine Glioma. Cancer Cell, 2017. 32(4): p. 520–537 e5.

53. Puget, S., et al., Mesenchymal transition and PDGFRA amplification/mutation are key distinct oncogenic events in pediatric diffuse intrinsic pontine gliomas. PLoS One, 2012. 7(2): p. e30313.

54. Thorslund, T., et al., Histone H1 couples initiation and amplification of ubiquitin signalling after DNA damage. Nature, 2015. 527(7578): p. 389–93.

55. Doil, C., et al., RNF168 binds and amplifies ubiquitin conjugates on damaged chromosomes to allow accumulation of repair proteins. Cell, 2009. 136(3): p. 435–46.

56. Kolas, N.K., et al., Orchestration of the DNA-damage response by the RNF8 ubiquitin ligase. Science, 2007. 318(5856): p. 1637–40.

57. Dos Santos Passos, C., et al., Design of genetically encoded sensors to detect nucleosome ubiquitination in live cells. J Cell Biol, 2021. 220(4).

58. Hu, Y., et al., RAP80-directed tuning of BRCA1 homologous recombination function at ionizing radiation-induced nuclear foci. Genes Dev, 2011. 25(7): p. 685–700.

59. Curtin, N.J. and C. Szabo, Poly(ADP-ribose) polymerase inhibition: past, present and future. Nat Rev Drug Discov, 2020. 19(10): p. 711–736.

60. Javle, M. and N.J. Curtin, The role of PARP in DNA repair and its therapeutic exploitation. Br J Cancer, 2011. 105(8): p. 1114–22.

61. Ausejo-Mauleon, I., et al., TIM-3 blockade in diffuse intrinsic pontine glioma models promotes tumor regression and antitumor immune memory. Cancer Cell, 2023. 41(11): p. 1911–1926 e8.

62. Muller, L., P. Aigner, and D. Stoiber, Type I Interferons and Natural Killer Cell Regulation in Cancer. Front Immunol, 2017. 8: p. 304.

63. Zitvogel, L., et al., Type I interferons in anticancer immunity. Nat Rev Immunol, 2015. 15(7): p. 405–14.

64. Le Bert, N., et al., STING-dependent cytosolic DNA sensor pathways regulate NKG2D ligand expression. Oncoimmunology, 2014. 3: p. e29259.

65. Zhang, Y., et al., Histone H3K27 methylation modulates the dynamics of FANCD2 on chromatin to facilitate NHEJ and genome stability. J Cell Sci, 2018. 131(12).

66. Werbrouck, C., et al., TP53 Pathway Alterations Drive Radioresistance in Diffuse Intrinsic Pontine Gliomas (DIPG). Clin Cancer Res, 2019. 25(22): p. 6788–6800.

67. Kim, H., et al., Targeting the ATR/CHK1 Axis with PARP Inhibition Results in Tumor Regression in BRCA-Mutant Ovarian Cancer Models. Clin Cancer Res, 2017. 23(12): p. 3097–3108.

68. Chornenkyy, Y., et al., Poly-ADP-Ribose Polymerase as a Therapeutic Target in Pediatric Diffuse Intrinsic Pontine Glioma and Pediatric High-Grade Astrocytoma. Mol Cancer Ther, 2015. 14(11): p. 2560–8.

69. Zarghooni, M., et al., Whole-genome profiling of pediatric diffuse intrinsic pontine gliomas highlights platelet-derived growth factor receptor alpha and poly (ADP-ribose) polymerase as potential therapeutic targets. J Clin Oncol, 2010. 28(8): p. 1337–44.

70. Baxter, P.A., et al., A phase I/II study of veliparib (ABT-888) with radiation and temozolomide in newly diagnosed diffuse pontine glioma: a Pediatric Brain Tumor Consortium study. Neuro Oncol, 2020. 22(6): p. 875–885.

71. Ngoi, N.Y.L., et al., Targeting the replication stress response through synthetic lethal strategies in cancer medicine. Trends Cancer, 2021. 7(10): p. 930–957.

72. Pommier, Y., M.J. O’Connor, and J. de Bono, Laying a trap to kill cancer cells: PARP inhibitors and their mechanisms of action. Sci Transl Med, 2016. 8(362): p. 362ps17.

73. Ronson, G.E., et al., PARP1 and PARP2 stabilise replication forks at base excision repair intermediates through Fbh1-dependent Rad51 regulation. Nat Commun, 2018. 9(1): p. 746.

74. Moreno-Lama, L., et al., Coordinated signals from PARP-1 and PARP-2 are required to establish a proper T cell immune response to breast tumors in mice. Oncogene, 2020. 39(13): p. 2835–2843.

75. Langelier, M.F., et al., Clinical PARP inhibitors allosterically induce PARP2 retention on DNA. Sci Adv, 2023. 9(12): p. eadf7175.

76. Larson, J.D., et al., Histone H3.3 K27M Accelerates Spontaneous Brainstem Glioma and Drives Restricted Changes in Bivalent Gene Expression. Cancer Cell, 2019. 35(1): p. 140–155 e7.

