## Supplementary figures and images for "H3K27M diffuse midline glioma is homologous recombination defective and sensitized to radiotherapy and NK cell-mediated antitumor immunity by PARP inhibition"

### Supplementary Figures

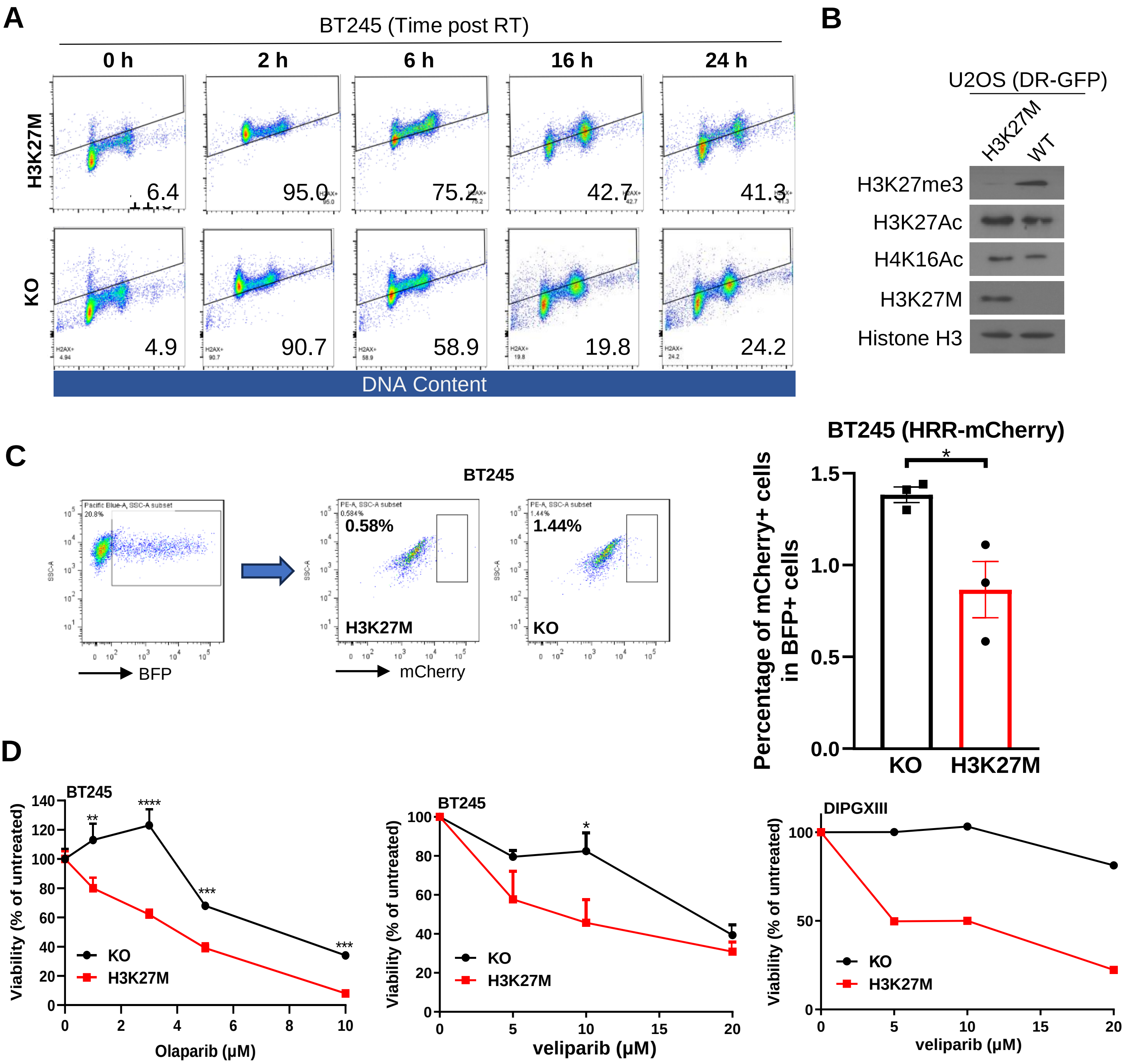

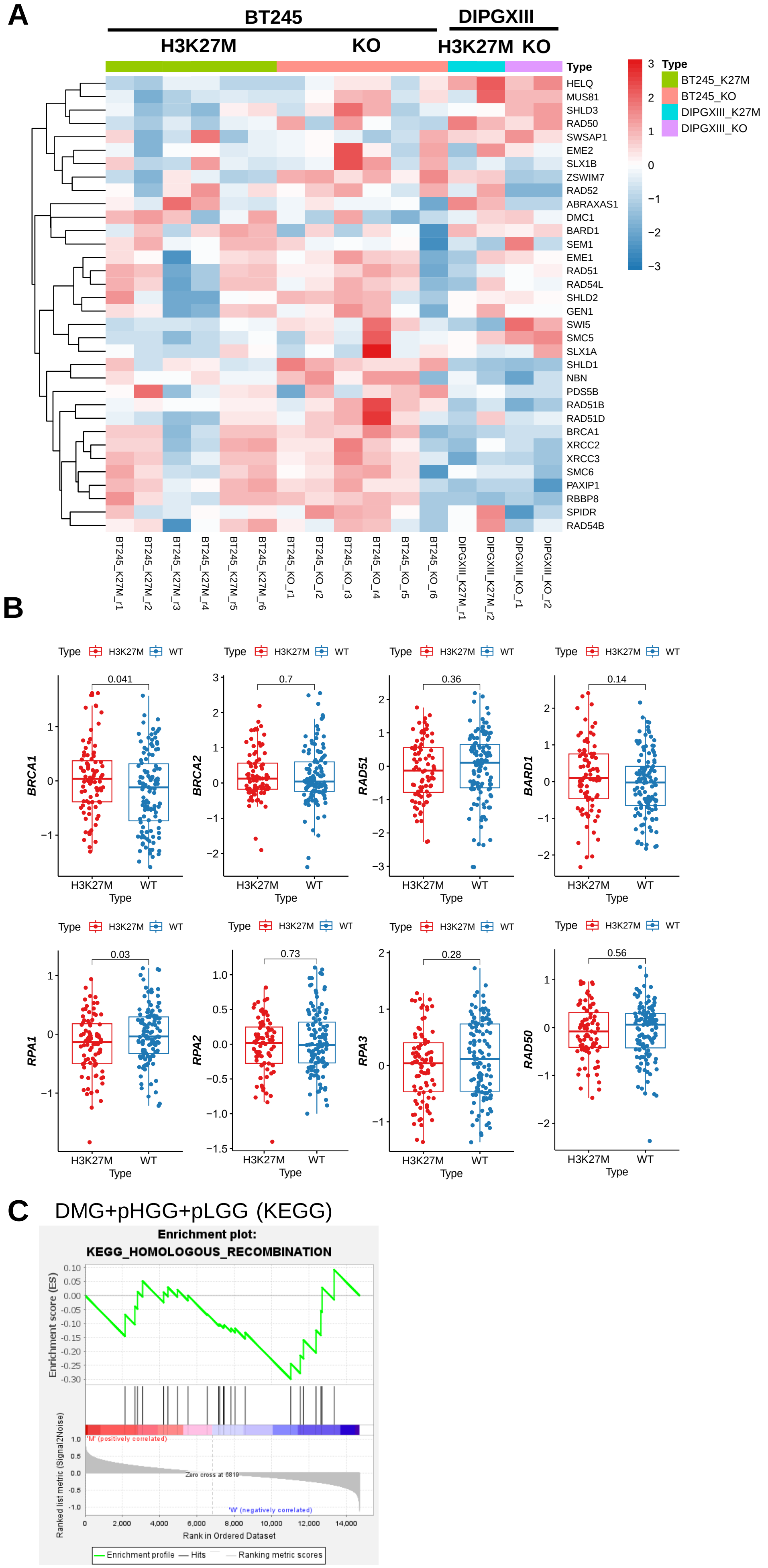

#### Figure S3

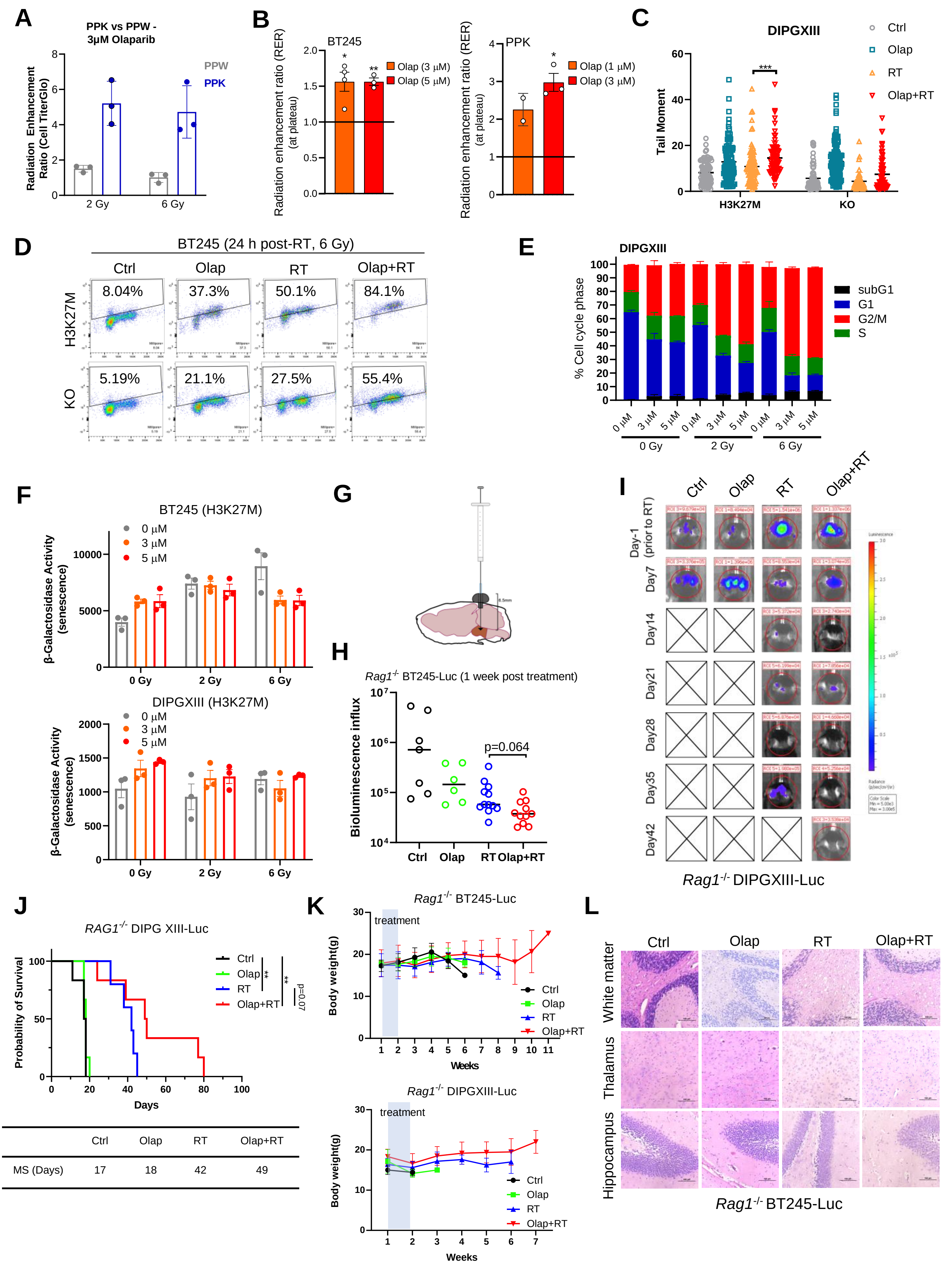


|  | Ctrl | Olap | RT | Olap+RT |
|---|---|---|---|---|
| MS (Days) | 17 | 18 | 42 | 49 |

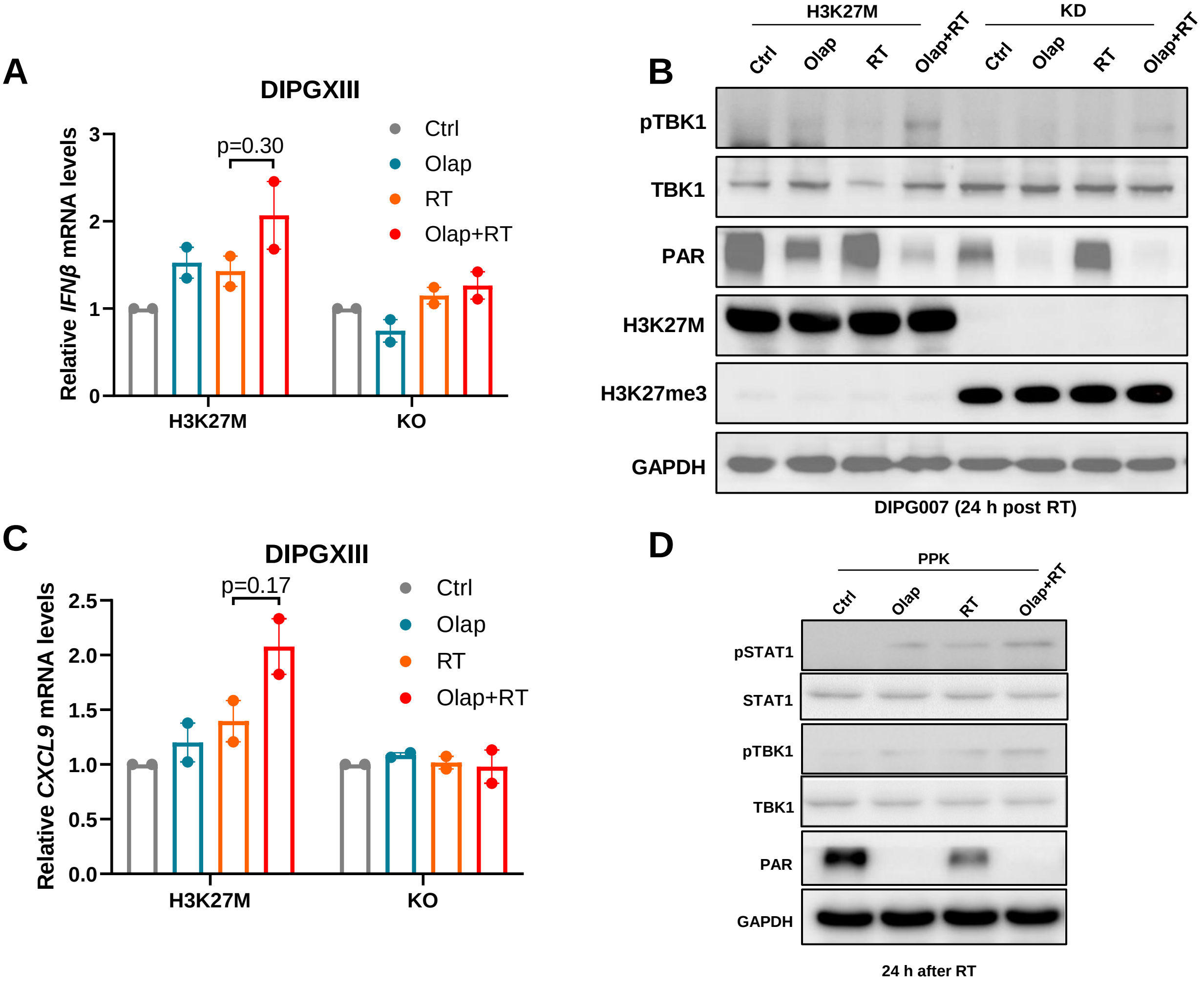

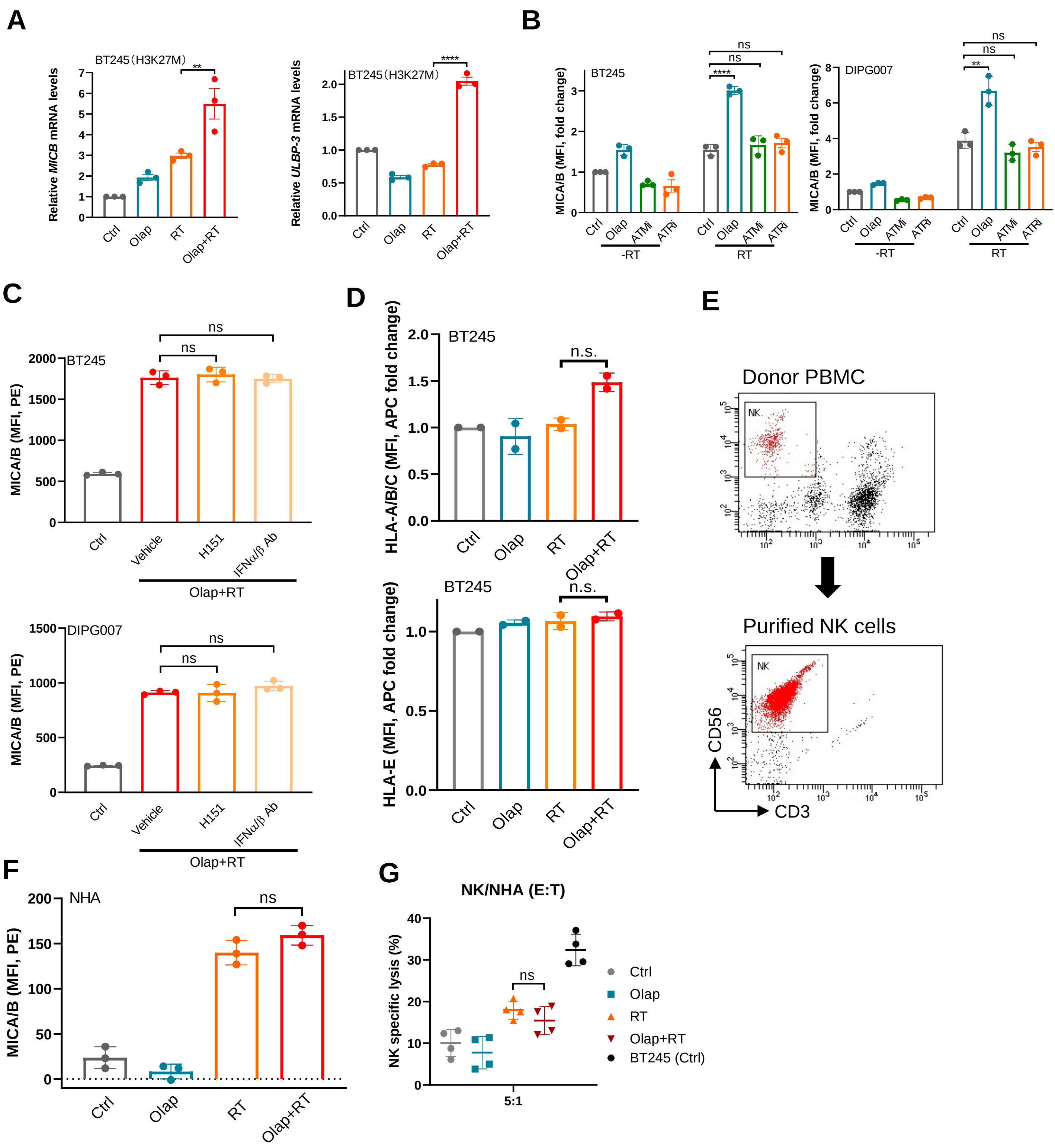

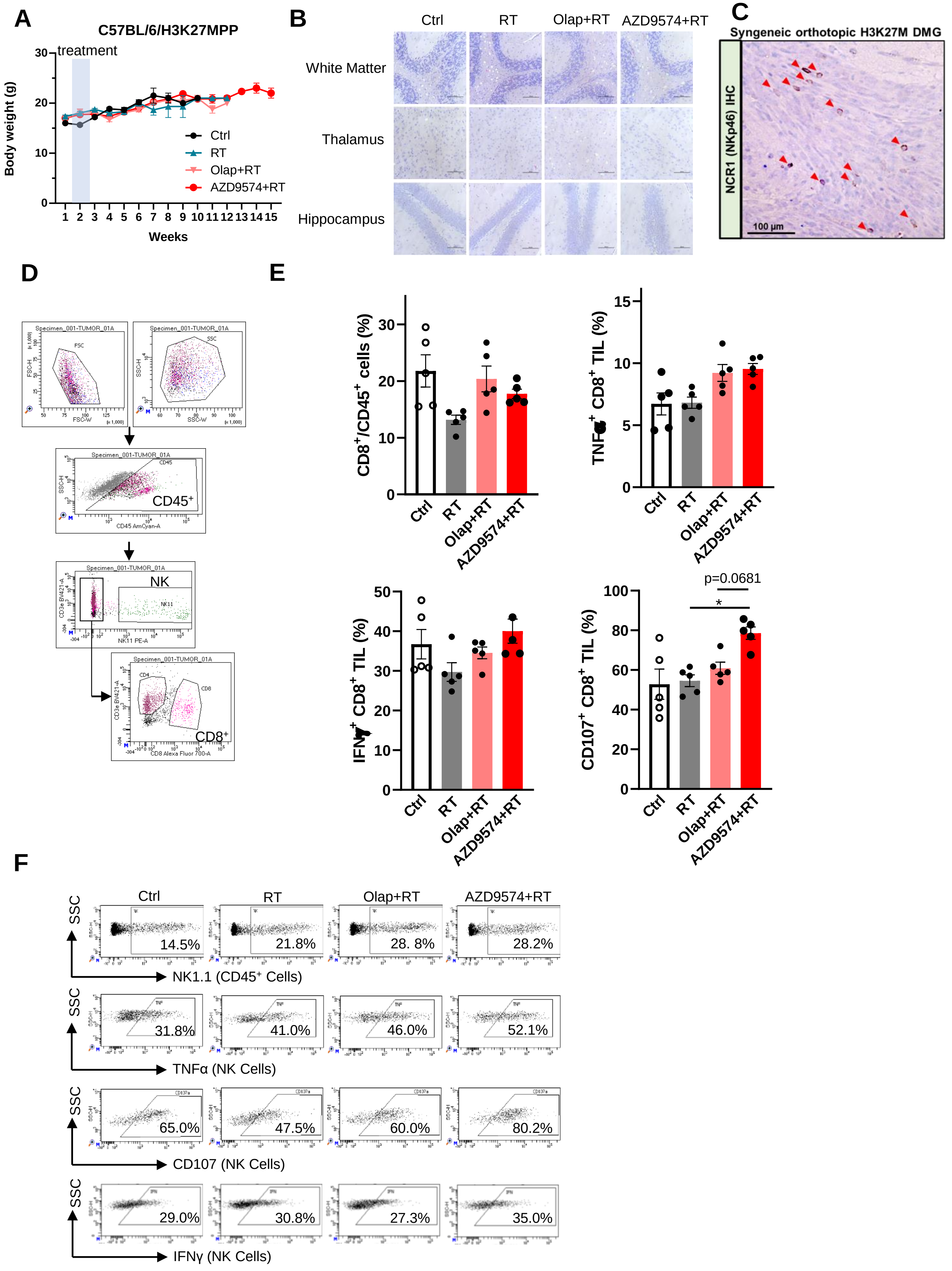
