## Supplementary Materials for "H3K27M diffuse midline glioma is homologous recombination defective and sensitized to radiotherapy and NK cell-mediated antitumor immunity by PARP inhibition"

Running title: Improving RT and NK cell immunity in H3K27M DMG.

* These authors contributed equally to this work

Corresponding authors: Qiang Zhang[†] and Meredith A Morgan[†], Department of Radiation Oncology, University of Michigan, Ann Arbor, Michigan, USA, 48109. Xinjun Wang Department of Neurosurgery, The Third Affiliated Hospital of Zhengzhou University, Zhengzhou, 45000, China.

[†]Lead contacts

### Supplementary Materials and Methods

**γH2AX flow assay.** BT245 and DIPGXIII (H3K27M and KO) cells were treated with accutase, washed with ice-cold PBS, and fixed at a concentration of 1 x $10^6$ cells/mL in ice-cold 70% ethanol. For γH2AX analysis, samples were incubated with a mouse γH2AX antibody (clone JBW301; Millipore) overnight at 4°C followed by incubation with a FITC-conjugated secondary antibody (Sigma), as previously described [1]. For quantification of γH2AX positivity, a gate was arbitrarily set on the control, untreated sample to define a region of positive staining for γH2AX of approximately 5%. This gate was then overlaid on the RT and/or olaparib-treated samples.

**Immunofluorescence analysis.** BT245 cells were seeded in 6-well plates and treated with radiation (2 Gy or 6 Gy). For RAD51 staining, cells were harvested after 24 h. For γH2AX, MDC1, RNF8, and FLAG-RNF168 staining, cells were harvested at 1 hr post-RT. Cells were then fixed for 15 min in ice-cold 4% paraformaldehyde, 2% sucrose, 0.5% Triton X-100 solution followed by permeabilization for 10 min in 0.5% Triton X-100/PBS. Cells were then stained with anti-RAD51 antibody (sc-8349; Santa Cruz), anti-γH2AX (JBW301, Millipore), anti-MDC1 (MA5-27650, ThermoFisher), anti-RNF8 (sc-271462, Santa Cruz Biotechnology) or anti-FLAG (M2, Sigma) and mounted with a drop (~10 µL) of ProLong™ Gold Antifade with DAPI (Invitrogen). Images were captured using an Olympus IX71 FluoView confocal microscope (Olympus America) with a 60x oil objective and Nikon NIS-Elements software. Images were then prepared using Fiji (NIH) software by equivalently adjusting only for brightness and contrast. At least 30-50 cells from each treatment condition were evaluated.

**Irradiation.** Irradiations for both cultured cells and tumor bearing animals were performed using a Philips RT250 (Kimtron Medical) at a dose rate of ~2 Gy/min in the University of Michigan, Rogel Cancer Center Experimental Irradiation Shared Resource. Dosimetry was performed using an ionization chamber connected to an electrometer system that is directly traceable to a National Institute of Standards and Technology calibration. For mouse brainstem irradiation, the upper part of brain and body were shielded with lead.

**HRR reporter assay.** For DR-GFP reporter, U2OS cells stably expressing a DR-GFP reporter plasmid [2] were infected with lentivirus expressing pInducer20/H3.3 WT or H3.3-K27M (generously provided by Dr. Efrat Shema, Weizmann Institute of Science, Israel) and cultured in DMEM medium with doxycycline (0.5 µg/mL). Cells were infected with adenovirus expressing the restriction enzyme I-SceI, which cleaves the defective DR-GFP gene cassette, while homology-directed repair of this break drives GFP expression. The extent of HR repair was then quantified by flow cytometric analysis of GFP expression. For measuring HRR in DMG cells, pLCN DSB Repair Reporter (DRR) (#98895, Addgene) was introduced into BT245 parental and KO cells by lentiviral infection. Stable clones were selected with neomycin selection (400 µg/mL) and further transfected with pCAGGS DRR mCherry Donor EF1a BFP (#98896, Addgene) and infected with I-SceI adenovirus as described in the previous study [3]. The transfection efficiency was controlled based on BFP expression from the donor cassette. The percentage of mCherry positive cells in the $BFP^+$ populations indicated the HRR efficiency.

**Cell titer glo assay.** BT245 and DIPGXIII cells were seeded in 6-well plates and treated with olaparib (0, 1, 3, 5, or 10 µM) or veliparib (0, 5, 10, or 20 µM). After 72 h, cells were harvested, counted, and seeded cells to opaque-walled 96-well plates. Cells were then cultured in incubator for 5 days. Cell viability was measured using the CellTiter-Glo luminescent cell assay kit (Cat.# G7571, Promega) according to the manufacture's instruction. Briefly, 96-well plates were put at room temperature for 15 min followed by adding 80 µL of Cell Titer-Glo® Reagent/well to induce cell lysis. Plates were then incubated at room temperature for 25 min to stabilize luminescent signal. Cell luminescence was measured by Cytation 3 plate reader.

**Neurosphere survival assay**. BT245 parental and PPK cells were plated into 6 well dishes ($2\times10^5$ cells/well). Forty-eight hours later, samples were treated with Olaparib (3 or 5 µM BT245 cells; 1 or 3 µM PPK cells), 1 h prior to RT. After 24 h, spheres were dissociated using Accutase (StemCell Technologies) and plated as single cell suspensions in quadruplicate in 96 well plates (Corning Costar #3610) which were transferred to a BioTek BioSpa 8 Automated Incubator (Agilent Technologies) and imaged every 12 h for 10-14 days with a BioTek Cytation 5 Cell Imaging Multimode Reader (Agilent). Images were processed for data acquisition using BioTek GenV software (Agilent). Each well was imaged at t0 to determine the exact number of cells seeded per well. Neurospheres with an area larger than 2600 $\mu m^2$ were counted as surviving spheres. Plating efficiency (PE), determined by the number of surviving spheres normalized to the number of cells at t0, was monitored over time for each condition, and the PE at the time a sample reached its growth plateau was used to calculate the surviving fraction (SF). Radiation survival was normalized for drug toxicity and the radiation enhancement ratio (RER) was calculated as the ratio of the radiation survival under control conditions divided by the radiation survival after drug exposure. A RER value significantly greater than 1 indicates radiosensitization.

**Western blot.** For whole cell protein extracts, cells were harvested and resupended in RIPA buffer (50 mM Tris [pH 7.5], 1% NP40, 0.5% SDS, 150 mM NaCl, 1 mM EDTA [pH 8.0]), supplemented with protease and phosphatase inhibitors (Roche). Protein concentration was determined using Bradford protein assay (Bio-Rad). Cell lysates with equal amount were denaturated in 4× Laemmli buffer for 10 minutes at 100°C heat block. Samples were then resolved by SDS-PAGE and transferred to PVDF membranes (0.2 μm). The antibodies against H3K27M (1:3000, #74829, Cell Signaling Technology), H3K27me3 (1:3000, #9733, Cell Signaling Technology), H3K27ac (1:3000, #8173, Cell Signaling Technology), H4K16ac (1:2000, #13534, Cell Signaling Technology), PAR (1:2000, 10H, ab14459, Abcam), PARP1 (1:1000, #9532, Cell Signaling Technology), PARG (1:1000, #66564, Cell Signaling Technology), H1.2 (1:1000, ab17677, Abcam), Polyubiquitin (K63-linkage-specific) (1:1000, BML-PW0600, Enzo), BRCA1 (1:1000, sc-6954, Santa Cruz), RAP80 (1:1000, #14466, Cell Signaling Technology), pTBK1 (1:500, #5483, Cell Signaling Technology), TBK1 (1:1000, #3013, Cell Signaling Technology), pSTAT1 (1:500, #9167, Cell Signaling Technology), STAT1 (1:1000, #14994, Cell Signaling Technology), and GAPDH (1:5000, #5174, Cell Signaling Technology).

**Dot blot.** For detecting PAR levels in the mouse brainstems, tissues were harvested at 1h and 4h after the last olaparib and RT treatment, and then smashed using a cryogenic tissue crusher on dry ice and resuspended in RIPA buffer with protease and phosphatase inhibitors. Protein concentration was determined using Bradford protein assay (Bio-Rad). Equal amount of tissue lysates (4 μg) was spotted onto a nitrocellulose membrane. After air drying, the membrane was blocked with 0.15 M NaCl, 0.01 M Tris–HCl, pH 7.4, 0.1% Tween 20 supplemented with 5% milk and extensively washed with the same buffer. The membrane was probed with anti-PAR (10H) antibody.

**Ubiquitination assay.** 293T cells expressing H3K27M or WT histone were transfected with His-ubiquitin plasmid by Lipofectamine 2000. After 48 h, cells were treated with RT (10 Gy) and washed with cold PBS for three times at 30 min post-RT. Cells were then lysed in denaturing buffer (6 M Guanidinium HCl, 100 mM $NaH_2PO_4$, 10 mM Tris-HCl, 5 mM β-mercaptoethanol, pH 8.0) and sonicated. Total cell extracts (2 mg) were then incubated with 100 μL of Ni-NTA Agarose (Qiagen) at room temperature for 4 h. The Ni-NTA beads were washed once with 1 mL of denaturing incubation buffer and three times with 1 mL of washing buffer (8 M urea, 100 mM $NaH_2PO_4$, 10 mM Tris-HCl, 20 mM imidazole, 5 mM β-mercaptoethanol, pH=6.3). Samples were resolved by SDS-PAGE for detecting histone H1.2 ubiquitination.

**Chromatin fractionation.** Parental and H3K27M KO BT245 cells were treated with RT (10 Gy) and lysed with buffer A (50 mM HEPES [pH 7.9], 10 mM KCl, 1.5 mM $MgCl_2$, 0.34 M sucrose, 10% glycerol, 1mM DTT, 0.1% Triton X-100, protease inhibitor cocktail) on ice for 20 min. After centrifugation (800 x g, 10min, 4°C), pellets including cell nuclei were further lysed with buffer B

(3 mM EDTA, 0.2 mM EGTA, 1 mM DTT, protease inhibitor cocktail). After second centrifugation (14,000 x g, 10 min, 4°C), pellets containing the chromatin fraction were washed and sonicated in RIPA lysis buffer for immunoblot analysis.

**Bioinformatics analysis.** Bulk RNA-seq data for HRR related gene expression analysis between parental H3K27M and KO BT245 and DIPGXIII cells were required from Dr. Nada Jabado and analyzed [4]. DMG/pHGG patient mRNA sequencing data including H3K27M (83 cases) and WT (118 cases) were downloaded from PedcBioportal [5]. mRNA data (53 DIPG samples) used for GSEA analysis (Supplementary Figure S2C) were accessed from UCSC Xena (cohort: Pediatric diffuse intrinsic pontine gliomas) [6]. Student's t-test was used when comparing two groups. All statistical analyses were performed using R (version 4.3.1). GSEA software (version 4.2.3) was used for KEGG analysis.

**RNA-seq data analysis.** BT245 cells treated with olaparib (3 µM pre-RT) and/or RT (8 Gy) were harvested at 72 h post-RT. Total mRNAs were extracted using RNeasy Mini Kit (Qiagen) and DNase digestion (Qiagen) and applied for bulk RNA sequencing at the Advanced Genomics Core of University of Michigan. RNA-seq data were deposited in the NCBI's Gene Expression Omnibus database (GSE268375). For data analysis, Fastq files were aligned and quantified by the Advanced Genomics Core. Differential gene expression (DGE) was run with DESeq2 in R (v4.3.3) for each comparison. Before DGE, genes that expressed less than 10 and had little to no variance were filtered out, and counts were normalized. Volcano plots were created using ggplot2. Gene set enrichment analysis (GSEA) was run with clusterProfiler using the Gene Ontology (GO) databases.

**Comet assay.** Single-cell gel electrophoretic comet assays of BT245 and DIPGXIII parental and isogenic KO cells were performed under alkaline conditions according to the previous protocol [7]. Briefly, cells were irradiated (6 Gy) with or without olaparib (3 µM, 1h pre-RT) and collected after 24 h. $2 \times 10^4$ cells/mL were combined with 1% Low Melting Point Agarose (Sigma) at 40°C and immediately pipetted onto slides (R&D Systems). For cellular lysis, the slides were immersed in alkaline lysis solution (1.2 M NaCl, 100 mM $Na_2EDTA$, 0.1% sodium lauryl sarcosinate, 0.26 M NaOH, pH > 13) overnight at 4°C in the dark, followed by washing in the rinse and electroporation buffer (0.03 M NaOH, 2 mM $Na_2EDTA$, pH ~12.3) for 30 min with two repeats. Then, the slides were subjected to electrophoresis at 20 V for 25 min and stained in 2.5 µg/mL propidium iodide for 15 min. All images were taken with a fluorescence microscope and analyzed by Comet Assay IV software (Perceptive Instruments).

**Cell apoptosis and senescence assay.** BT245 and DIPGXIII cells were treated with olaparib (0, 3, or 5 µM, 1 hr pre-RT) and radiation (0, 2, 6 Gy). After 72 h, cell apoptosis and senescence were monitored using an Annexin V-FITC/7-AAD staining kit (BioLegend, #640922) and Senescence β-Galactosidase Staining Kit (Cell Signaling Technology, #9860) according to the manufactures' instructions, respectively.

***IFNβ* promoter-GFP reporter assay.** The pLKO.1-hygro-*IFNβ*–GFP reporter construct generously provided by Dr. Roger A. Greenberg [8] was transfected into 293T cells with lentiviral packaging plasmids. BT245 and DIPG007 cells were infected and selected with 50 µg/mL hygromycin for 2 weeks. The BT245 (H3K27M and KO)-*IFNβ*–GFP and DIPG007 (H3K27M and KO)-*IFNβ*–GFP reporter cells were treated with olaparib (3 µM, 1 h pre-RT) and/or radiation (8 Gy) and harvested after 3 days. GFP expression levels of reporter cells with various treatment were measured by flow cytometry (BD Biosciences). The change in median fluorescence intensity (MFI) for indicated treatments was obtained by subtraction of background GFP levels [9].

**qPCR.** RNA was isolated by RNeasy Mini Kit (Qiagen) and DNase digestion (Qiagen) from cells with indicated treatment. RNA concentration and purity were measured using a Nanodrop spectrophotometer (Thermo Fisher Scientific). cDNA was reverse-transcribed using High-Capacity RNA-to-cDNA Kit (Applied Biosystems). Relative indicated gene expression levels were determined by quantitative PCR (qPCR) using Fast SYBR® Green Master Mix (Invitrogen) on a StepOnePlus™ Real-Time PCR System (Thermo Fisher Scientific) and fold change (ΔΔCt) method normalized to GAPDH. The following qPCR primers were used: human *IFNβ*, 5′-ATGACCAACAAGTGTCTCCTCC-3′, 5′-GCTCATGGAAAGAGCTGTAGTG-3′; human *CXCL9*, 5′-GTGGTGTTCTTTTCCTCTTGGG-3′, 5′-ACAGCGACCCTTTCTCACTAC-3′; human *CXCL10*, 5′-CTCCAGTCTCAGCACCATGA-3′, 5′-GCTCCCCTCTGGTTTTAAGG-3′; human *MICA*, 5'-CCTTGGCCATGAACGTCAGG-3', 5'-CCTCTGAGGCCTCGCTGCG-3', human *MICB*, 5'-ACCTTGGCTATGAACGTCACA-3', 5'-CCCTCTGAGACCTCGCTGCA-3', human *ULBP-3*, 5'-CTCGCGATTCTTCCGTACCT-3', 5'-TCTGGACCTCACACCACTGT-3', mouse *IFNβ*, 5'-CCCTATGGAGATGACGGAGA3', 5'-CTGTCTGCTGGTGGAGTTCA-3' mouse *Cxcl9*, 5′-CCTAGTGATAAGGAATGCACGATG-3′, 5′-CTAGGCAGGTTTGATCTCCGTTC-3′; and mouse *Cxcl10*, 5′-CCTGCCCACGTGTTGAGAT-3′, 5′-TGATGGTCTTAGATTCCGGATTC-3′.

**Flow cytometry.** To analyze cell surface MICA/B, ULBP-3, Rae1, and H60 expression, human BT245, DIPG007, and mouse H3K27MPP cells with indicated treatment were incubated with accutase to generate single-cell suspension in cell staining buffer (2% FBS in $Ca^{2+}/Mg^{2+}$ free free PBS). Cells were then incubated with PE- or FITC-conjugated anti-human PD-L1 antibody for 1 h at room temperature in the dark. Stained cells were washed in the staining buffer and analyzed by flow cytometry (BD Biosciences). IgG antibody with no specific binding was used as a background MFI. The NKG2D ligand expression levels on the cell surface were analyzed in FlowJo 10 software. For the analysis of brainstem TME in the syngeneic orthotopic DMG mice, tissues (n=5 per group) were cut into small pieces and transferred in 50 mL tubes containing 10 mL digestion buffer (3 mg/mL Collagenase IV, Worthington #LS004209, 1 mg/mL DNase I, Worthington #LS002007, and 2 mg/mL Soybean Trypsin Inhibitor, Worthington #LS003587 in PBS), and incubated at 37℃ shaker (180 rpm) for 30 minutes. Digested tissues were further washed with PBS and passed through a 70µm mesh cell strainer followed by removal of erythrocytes, myelin, and cell debris using ACK buffer (#A1049201, Gibco) and myelin removal beads (Miltenyi #130-096-731) and columns (Miltenyi #130-042-201) [10]. Cells were then treated with PMA (5 ng/mL, P1585, Sigma), Ionomycin (500 ng/mL, I9657, Sigma,), GolgiPlug (1:1,000, #555029, BD Biosciences,) and GolgiStop (1:1,000, #554724, BD Biosciences) in RPMI in 37˚C incubator for 4 h. After treatment, cells were stained in FACS buffer for surface markers and then fixed/permeabilized by using a fixation/permeabilization kit (00-5123-43, 00-5223-56, eBiosciences). After washing with permeabilization buffer, cells were stained with the cytokines markers and added with counting beads (C36995, Invitrogen). Flow cytometry analysis was performed using the BD Fortessa instrument and FlowJo 10 software. Antibodies used for staining include: anti-mouse CD45 (V450, 30-F11, #560501, BD Biosciences), anti-mouse NK1.1 (PE, PK136, #108708, BioLegend), anti-mouse CD8a (APC-R700, 53-6.7, #56-0081-82, eBioscience), anti-mouse IFNγ (BV786, XMG1.2, #563773, BD Biosciences), anti-mouse TNFα (BV605, MP6-XT22, #506329, BioLegend), and anti-mouse CD107 (PE-Cy7, 1D4B, #560647, BD Biosciences).

**NK cell killing assay.** NK92 cells from ATCC were cultured in MyeloCult H5100 (#05150, StemCell Technologies, 12.5% horse serum, and 20 ng/mL IL-2). Primary human NK cells were isolated from a healthy donor using the EasySep™ Direct Human NK Cell Isolation Kit (Catalog # 19665, STEMCELL) and expanded in NK MACS medium containing NK MACS Supplement (1:100, #130-114-429, Miltenyi), 5% AB serum (H5667, Sigma), IL-2 (500 IU/mL), and IL-15 (140 IU/mL). For NK cell lysis assay, BT245 and DIPG007 cells were treated with olaparib (3 µM) and/or RT (8 Gy) for 72 h and then fluorescently labeled with calcein-AM dye. After washing with PBS for three times, cells were incubated with either NK92 or ex vivo expanded and activated primary NK cells at two different effector to target (effector:target, E:T) ratios (5:1 and 10:1) for 4-

6 h. Calcein-AM release was measured using Cytation 3 plate reader (excitation 485 nm and emission 530 nm). The percentage of calcein release which indicated lysis of target cell was calculated as previously reported [11].

**In vivo orthotopic DMG model and treatment**. An implantable guide-screw system was used to generate orthotopic brainstem DMG models as previously described [12]. *Rag1*$^{-/-}$ or C57BL/6 mice were anesthetized and simultaneously administrated with carprofen analgesia followed by removing scalp fur and sterilizing the surgical site. A longitudinal incision was made to expose the skull, and the entry point 1.0 mm right to lambda and just posterior (0.8 mm) to lambdoid suture was made in the skull using a handheld. Approximately $2\times10^5$ cells in a volume of 2μL were implanted into the brainstem at a depth of 6.5 mm with a 10μL Hamilton syringe (Supplementary Figure S3G). A PHD Ultra multi-syringe pump (Harvard Apparatus) was used to facilitate simultaneous implantation of up to 8 mice at a time. After injections, mice were sutured and treated with triple antibiotic ointment. Atipamezole was administered to reverse anesthesia on a heating pad. Mice were then fed with diet gel and monitored daily for 10 days, including a second dose of carprofen one day after surgery.

PARP inhibitors olaparib and AZD9574 were synthesized and provided by AstraZeneca with an agreement. Olaparib was dissolved in dimethyl sulfoxide (Sigma) and then diluted in 10% 2-hydroxypropyl-β-cyclodextrin (Cayman Chemicals), and AZD9574 was dissolved in 1M methanesulfonic acid (MSA) (Sigma) then diluted in sterile dd$H_2O$ followed by adjusting pH value to 3-3.2 using 1M NaOH. Olaparib (100 mg/kg) or AZD9574 (3 mg/kg) was administered by oral gavage 1 h before radiation and continued for 6 days (Scheduled timelines as shown in Figure 3F and 6A). Mice were then sedated using 2.5% isoflurane and treated with RT (2Gy, per fraction) on days 0-2, 4-6. Moribund mice were euthanized, and survival data was analyzed using the Kaplan-Meier method by GraphPad Prism 10.0. For the *Rag1*$^{-/-}$/BT245-Luc mice, Bioluminescence imaging (BLI) was measured once weekly by administering sterile-filtered 30 mg/mL luciferin solution (Syd Laboratories) via IP injection. BLI signal was measured by IVIS Spectrum imaging station and Luminescent Flux Values were obtained for each tumor. BLI flux values were used to randomize mice into four treatment groups (6-12 mice/group) based on average BLI flux signal. For NK1.1 depletion, anti-mouse NK1.1 blocking antibody (BioXCell #BE0036) or IgG2a isotype control (BioXCell #BE0085) were concurrently administered with olaparib and RT by intraperitoneally injecting antibody (200 μg/mouse) every 3 days starting at day -1 (day 0 represents the start of radiation treatment) for a total of 4 doses (schedule shown in Figure 6D).

**Immunohistochemistry staining.** DMG tumor bearing mouse brains were harvested and fixed in 4% paraformaldehyde, and paraffin-embedded blocks were cut into 5 μm sections. Slides were de-waxed in xylene followed by rehydration in a standard alcohol series. Antigen retrieval process was performed by using microwave heating in a retrieval buffer (Vector Labs #H-3301), followed by blocking of endogenous peroxidase in 3% $H_2O_2$. The H3K27M (1:200, #3386474, Millipore Sigma) and NCR1 (1:200, ab233558, Abcam) antibodies were added and incubated overnight at 4 °C. Slides were then applied with a secondary-HRP labeled mouse or rabbit antibody detection system (Vector Labs #PK-6101, #PK-6102) followed by addition of 3,3'-diaminobenzidine (DAB) chromagen (Vector Labs #SK-4100). Slides were finally counter-stained with hematoxylin (Fisher Scientific) and dehydrated in 70, 95 and 100% ethanol and xylene. Images were acquired with an Olympus BX-51 microscope, Olympus DP71 digital camera, and DP Controller software.

### Supplementary Figure Legends

**Supplementary Figure 1**. **H3K27M causes a homologous recombination repair defect.** (**A**) H3K27M isogenic BT245 cells were assessed for γH2AX positivity by flow cytometry at the indicated times post-RT (6 Gy). (**B**) Western blot analysis of U2OS (DR-GFP) cells expressing either H3K27M or WT histone H3 with the indicated antibodies. Histone H3 represent loading controls. (**C**) Parental and H3K27M KO BT245 cells with pLCN DSB Repair Reporter (DRR) were transfected with HR donor (DRR mCherry Donor EF1a BFP cassette) and infected with I-SceI adenovirus. After 48 h, cells were harvested for flow cytometry analysis. Percentages of $mCherry^+$ cells, gated on BFP positive populations represented HRR efficiency. (**D**) CellTiter Glo analysis of BT245 and DIPGXIII parental and H3K27M KO cells with indicated concentrations of olaparib or veliparib treatment. Data represent 1 or 2 independent experiments.

**Supplementary Figure 2. H3K27M mutation abrogates RT-induced K63-linked polyubiquitination of histone H1 and recruitment of HRR proteins.** (**A**) Heatmap analysis showing the relative expression level of key HRR-related genes in BT245 and DIPGXIII H3K27M and isogenic KO cells. (**B**) Relative mRNA levels of HRR-related genes in H3K27M and WT DMG/pHGG patients from PedcBioPortal database (including 83 cases of H3K27M and 118 cases of WT) (**C**) KEGG-based GSEA analysis showed that HRR signaling pathway was not significantly enriched in H3K27M patients ($|NES| < 1$, p-value $> 0.05$, FDR $> 0.25$).

**Supplementary Figure 3. Selective radiosensitization of H3K27M DMG cells and orthotopic tumors by PARP inhibitor.** (**A**) CellTiter Glo analysis of PPK (H3K27M) and PPW (H3WT) cell viability after olaparib (3 µM) and RT (2 Gy or 6 Gy) treatment. (**B**) Neurosphere survival analysis of BT245 and PPK cells with RT (6 Gy) and/or olaparib. The radiation enhancement ratios (RER) were calculated by dividing survival fraction of olap+RT/olap into survival fraction of RT/untreated control. Values significantly >1 indicate radiosensitization. See the methods. (**C**) Alkaline comet assay to assess DNA damage levels in H3K27M and isogenic KO DIPGXIII cells in response to radiation (6 Gy) and/or olaparib (3 µM) treatment. (**D**) Representative results of γH2AX positive BT245 H3K27M and KO cells at 24 h after olaparib (3 µM) and RT (6 Gy) treatment. (**E**) Cell cycle distribution assessed by flow cytometry in DIPGXIII cells with after treatment with radiation and olaparib. (**F**) Percentage of senescent cells assessed by β-gal flow cytometry analysis in BT245 and DIPGXIII cells with after treatment with radiation and olaparib. (**G**) Schematic drawing of screw guide implantation of BT245-Luciferase cells into mouse pons. (**H**) Bioluminescent intensity in BT245-Luciferase tumors at 1 week after treatment. Data are presented as the mean ± SEM. (**I**) Representative in vivo bioluminescence images of the DIPGXIII-Luc tumors before, after, and during the treatment of olaparib and/or RT. (**J**) Survival analysis of *Rag1*$^{-/-}$ mice harboring DIPGXIII-Luc tumors treated as indicated (Ctrl n=6, olap n=6, RT n=5, RT+olap n=6). Data were analyzed using the log-rank test. (**K**) Mouse body weight before, after, and during the treatment of olaparib and/or RT for *Rag1*$^{-/-}$/BT245-Luc and *Rag1*$^{-/-}$/BT245-Luc study animals. Data are presented as the mean ± SEM. (**L**) Representative H&E staining images of white matter, thalamus, and hippocampus in the mouse brains after treatment with radiation and/or olaparib (scale bar: 100 µm).

**Supplementary Figure 4. PARP inhibitor and RT preferentially induce type I interferon signaling in H3K27M DMG cells.** (**A**) qPCR for *IFNβ1* mRNA in H3K27M and KO DIPGXIII cells with indicated treatment. (**B**) Western blotting analysis of pTBK1 and TBK1 levels in H3K27M and KO BT245 cells at 24 hr in response to olaparib and/or RT treatment. (**C**) qPCR analysis of mRNA levels of interferon-stimulated genes *CXCL9* in DIPGXIII cells with or without H3K27M mutation at 72 h after olaparib and/or RT treatment. (**D**) Western blotting analysis of pTBK1, TBK1, pSTAT1, and STAT1 levels in H3K27M mouse DMG cells at 24 hr in response to olaparib and/or RT treatment.

**Supplementary Figure 5. PARP inhibitor and RT increase expression of NKG2D activating ligands and induce NK cell lysis.** (**A**) qPCR for *MICB* and *ULBP-3* mRNA levels in BT245 cells at 72 h after olaparib and/or RT treatment. (**B**) Cell surface MICA/B levels of H3K27M parental

BT245 and DIPG007 cells in response to RT and/or indicated DDR inhibitor (olaparib 3 µM, AZD1390 50 nM, AZD6738 200 nM) treatment. (**C**) Flow cytometry analysis of cell surface MICA/B protein levels on BT245 and DIPG007 cells at 72 h following indicated treatment (olaparib 3 µM, H151 1 µM, IFNα/β neutralization antibody, 100 µg/mL). (**D**) Cell surface HLA-E and HLA-A/B/C levels of BT245 cells at 72 h following treatment of olaparib (3 µM) and/or RT (8 Gy). (**E**) Isolation and enrichment of primary human NK cells ($CD56^+/CD3^-$) from human donor whole blood. (**F**) Flow cytometry analysis of cell surface MICA/B in NHA cells at 72 h following olaparib (3 µM) and/or RT (8 Gy) treatment. (**G**) NK cell lysis of NHA cells co-cultured with primary activated NK cells after 72 h pretreatment of NHA cells with olaparib (3 µM) and/or RT (8 Gy).

**Supplementary Figure 6. PARP inhibitors and RT induce a therapeutic response in immune competent mice that is dependent upon NK cells.** (**A**) Mouse body weight before, after, and during the indicated treatment for C57BL/6/H3K27MPP DMG study animals. Data are presented as the mean ± SEM. (**B**) Images of H&E staining of white matter, thalamus, and hippocampus in the mouse brains after treatment with radiation, olaparib+RT, and AZD9574+RT (scale bar: 100 µm). (**C**) Immunohistochemistry staining of the murine NK cell marker NCR1 in C57BL/6/H3K27MPP DMG tumor. Scale bar, 100 µm. (**D**) Gating strategy to determine $CD45^+$, $NK1.1^+$, and $CD8^+$ cells in the mouse DMG tumors treated with RT with or without olaparib or AZD9574 as indicated in Figure 6A. (**E**) Flow cytometry analysis of the percentages of intratumoral $CD8^+$ cells (in the $CD45^+$ cell populations), and TNF-$\alpha^+$, $CD107^+$, and IFN$\gamma^+$ $CD8^+$ cells from mouse H3K27MPP DMG tumors treated as indicated. Data are the mean ± SEM (n = 5 mice/group). (**F**) Data are representative flow cytometry illustrations of the percentages of intratumoral NK cells (in the $CD45^+$ cell populations), and TNF-$\alpha^+$, $CD107^+$ NK cells in Figure 6E.

### References


1. Zhang, Q., et al., *FBXW7 Facilitates Nonhomologous End-Joining via K63-Linked Polyubiquitylation of XRCC4.* Mol Cell, 2016. **61**(3): p. 419-433.
2. Pierce, A.J., et al., *XRCC3 promotes homology-directed repair of DNA damage in mammalian cells.* Genes Dev, 1999. **13**(20): p. 2633-8.
3. Arnoult, N., et al., *Regulation of DNA repair pathway choice in S and G2 phases by the NHEJ inhibitor CYREN.* Nature, 2017. **549**(7673): p. 548-552.
4. Harutyunyan, A.S., et al., *H3K27M induces defective chromatin spread of PRC2-mediated repressive H3K27me2/me3 and is essential for glioma tumorigenesis.* Nat Commun, 2019. **10**(1): p. 1262.
5. Mackay, A., et al., *Integrated Molecular Meta-Analysis of 1,000 Pediatric High-Grade and Diffuse Intrinsic Pontine Glioma.* Cancer Cell, 2017. **32**(4): p. 520-537 e5.

6. Puget, S., et al., *Mesenchymal transition and PDGFRA amplification/mutation are key distinct oncogenic events in pediatric diffuse intrinsic pontine gliomas.* PLoS One, 2012. **7**(2): p. e30313.
7. Olive, P.L. and J.P. Banath, *The comet assay: a method to measure DNA damage in individual cells.* Nat Protoc, 2006. **1**(1): p. 23-9.
8. Harding, S.M., et al., *Mitotic progression following DNA damage enables pattern recognition within micronuclei.* Nature, 2017. **548**(7668): p. 466-470.
9. Zhang, Q., et al., *Potentiating the radiation-induced type I interferon antitumoral immune response by ATM inhibition in pancreatic cancer.* JCI Insight, 2024. **9**(6).
10. du Chatinier, A., et al., *Generation of immunocompetent syngeneic allograft mouse models for pediatric diffuse midline glioma.* Neurooncol Adv, 2022. **4**(1): p. vdac079.
11. Zhou, Z., et al., *Granzyme A from cytotoxic lymphocytes cleaves GSDMB to trigger pyroptosis in target cells.* Science, 2020. **368**(6494).
12. Marigil, M., et al., *Development of a DIPG Orthotopic Model in Mice Using an Implantable Guide-Screw System.* PLoS One, 2017. **12**(1): p. e0170501.
